# CRlSPR/Cas9 screening revealed *BlRC6-AS1*/BlRC6 mediates abiraterone resistance via NHEJ pathway-dependent A20 degradation in prostate cancer

**DOI:** 10.1101/2025.10.01.679907

**Authors:** Lan Li, Xue-Ni An, Yong-Tong Ruan, Rong Yang, Peng Li, Xiao-Yu Wu, Xian-Chao Huang, Ping Gao, Xiao-Ming Dong

## Abstract

Abiraterone acetate serves as the first-line therapeutic agent for prostate cancer (PCa) treatment. However, drug resistance frequently emerges. Employing a genome-wide CRISPR/Cas9 library screening strategy, we identified 523 long non-coding RNAs (lncRNAs) and 2183 protein-coding genes associated with abiraterone resistance. Notably, a pair of sense-antisense genes, *BIRC6-AS1*/*BIRC6*, was demonstrated to be a pivotal driver for abiraterone resistance. *BIRC6-AS1* depletion led to a reduction in both the mRNA and protein levels of BIRC6. Moreover, depletion of either *BIRC6-AS1* or BIRC6 enhanced the chemosensitivity of PCa cells to abiraterone both in vitro and in vivo settings. Further investigation revealed that *BIRC6-AS1* stabilized the mRNA of *BIRC6* through interaction with ILF2. Diminishing either *BIRC6-AS1* or BIRC6 predominantly suppressed non-homologous end joining (NHEJ) repair activity, resulting in the disassembly of 53BP1 foci at DNA damage sites and an increased accumulation of DNA damage in PCa cells induced by abiraterone. Mechanistically, BIRC6 interacted with A20 and facilitated the K48-linked ubiquitination and subsequent degradation of A20 at the K337 residue. Additionally, A20 knockdown effectively reversed the abiraterone sensitivity induced by *BIRC6-AS1* depletion. Collectively, we conducted a comprehensive screen to identify lncRNAs and protein-coding genes associated with abiraterone resistance and proposed that targeting *BIRC6-AS1*/BIRC6 axis represents a promising strategy to overcome abiraterone resistance in prostate cancer.

## Introduction

Primary prostate cancer (PCa) is an androgen-dependent cancer, relying on endogenous ligands to activate the androgen receptor (AR). It is initially responsive to androgen deprivation therapy (ADT), but most patients eventually progress to castration-resistant prostate cancer (CRPC). Abiraterone acetate, a first-line therapeutic agent targeting the steroidogenic enzyme CYP17A1, has been demonstrated to significantly prolong overall survival in men with CRPC ^1^. However, not all patients display a clinical response to abiraterone treatment, and even among those who initially achieve therapeutic efficacy, resistance may emerge over time. Despite the emergence of numerous hypotheses, including AR alterations, germline variants, and mechanisms associated with glucocorticoid ^2^, the genetic and molecular mechanisms underlying abiraterone resistance have not yet been fully elucidated.

To date, research endeavors dedicated to deciphering the mechanisms underlying abiraterone resistance have largely focused on protein-coding genes, whereas studies exploring the potential association between non-coding RNAs and abiraterone resistance are comparatively limited. Among the non-coding RNAs, long non-coding RNAs (lncRNAs) represent a substantial class that has been demonstrated to be implicated in chemotherapy resistance across various cancer types^3–6^. One of the emerging roles of lncRNAs lies in the regulation of protein - coding gene transcription, either *in cis* or *in trans*^7^. This implies the potential that lncRNAs may co-modulate drug resistance in conjunction with protein-coding genes. Thus, conducting a genome-scale functional characterization of both lncRNAs and protein -coding genes is critical for the identification of novel genes and underlying mechanisms that are implicated in chemotherapy resistance ^8^.

The genome-wide CRISPR-Cas9 knockout library screening approach empowers us to perform an impartial, high-throughput screening of drug resistance genes^9–13^. In this study, we performed a genome-wide CRISPR-based screen to characterize lncRNAs and protein-coding genes associated with abiraterone resistance in PCa cells. The results identified 523 lncRNAs and 2,183 protein-coding genes potentially implicated in conferring abiraterone resistance. Notably, four pairs of sense-antisense genes functionally modulated PCa cell resistance to abiraterone, among which the *BIRC6-AS1*/BIRC6 pair was selected for further analysis. Depletion of either *BIRC6-AS1* or BIRC6 enhanced cellular sensitivity to abiraterone *in vitro* and *in vivo*. Mechanistically, *BIRC6-AS1* stabilized *BIRC6* mRNA via interaction with ILF2, while BIRC6 knockout exacerbated abiraterone-induced DNA damage by inhibiting NHEJ repair activity, due to reduced ubiquitination and degradation of A20. These findings suggested that targeting the *BIRC6-AS1*/BIRC6 axis represents a promising strategy to overcome abiraterone resistance in PCa.

## Results

### CRISPR library screening identified *BIRC6-AS1*/*BIRC6* as a critical pair of genes for abiraterone resistance

It is well established that AR-negative PCa cell lines, including PC3 and DU145, exhibit resistance to androgen deprivation therapy and represent advanced disease stages. Although these cell lines remain sensitive to abiraterone, they demonstrate higher drug tolerance compared to AR-positive counterparts such as LNCaP cells. To systematically identify genome-wide factors contributing to abiraterone resistance, we performed CRISPR knockout (KO) library screening targeting both lncRNAs and protein-coding genes in PC3 cells, with subsequent validation in both PC3 and Du145 cells. The human lncRNA splicing-targeting CRISPR library^14^, consisting of 126,773 sgRNAs, was employed to target 10,996 lncRNAs. Concurrently, the human genome-wide lentiviral CRISPR sgRNA library^15^, containing 90,230 sgRNAs, was applied to target 18,424 protein-coding genes. These libraries were independently transduced into Cas9-expressing PC3 cells, which exhibited high cleavage efficiency (**Figures S1A-1C**) at a low multiplicity of infection (MOI) (∼0.3), followed by puromycin selection. The cells were subsequently treated with abiraterone or DMSO for 28 days and then harvested for high-throughput sequencing (**Figure 1A**).

**Figure 1.**
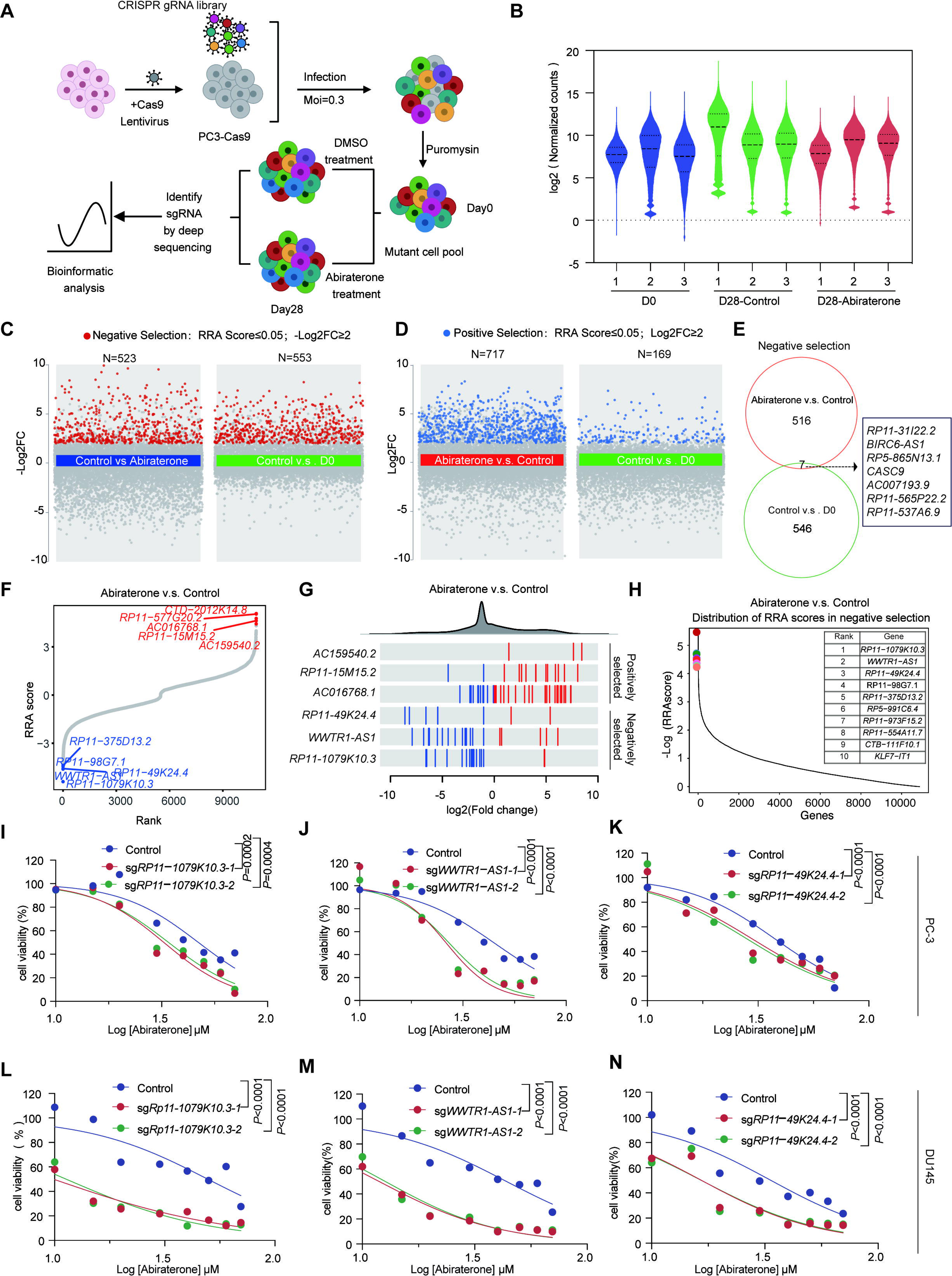
CRISPR library screening identified critical LncRNAs for abiraterone resistance. (**A**) Schematic of CRISPR-Cas9 screens: A lentiviral sgRNA library was transduced into PC3-Cas9 cells, which were then treated with DMSO or Abiraterone, respectively. After 28 days, sgRNAs were extracted for NGS. (**B**) Box plots displaying sgRNA distribution in the experimental groups from lncRNA CRISPR-Cas9 library: D0-DMSO (baseline), D28-DMSO (vehicle control), and D28-Abiraterone (treatment). (**C** and **D**) Volcano plots showing depleted (red; RRA Score ≤ 0.05, -log□FC ≥ 2) and enriched (blue; RRA Score ≤ 0.05, log□FC ≥ 2) genes. Screening analysis was performed with MaGeCK RRA. (**C**) Negative selection identified 523 abiraterone resistance-associated LncRNAs and 553 essential LncRNAs. (**D**) Positive selection revealed 717 LncRNAs associated with abiraterone sensitivity and 169 essential LncRNAs. (**E**) Venn diagram showed negatively selected genes from two comparisons: Abiraterone vs Control and Control vs D0. (**F**) MAGeCK analysis results displayed a ranking of genes based on their RRA scores. (**G**) Frequency distribution of log2 fold change for all sgRNAs (top) and log2 fold change of individual sgRNAs for representative candidates (bottom). Enriched and depleted sgRNA hits were indicated by red and blue vertical bars, respectively. (**H**) The RRA score distribution plot revealed the top 10 candidate LncRNAs associated with abiraterone resistance. (**I-N**) Cell viability assays in PC3 (**I-K**) and DU145 (**L-N**) cells treated with 0-70 μM abiraterone for 48h, following transduction with either control sgRNAs or sgRNAs targeting candidate lncRNAs: *RP11-1079K10.3* (**I** and **L**), *WWTR1-AS1* (**J** and **M**), and *RP11-49K24.4* (**K** and **N**). Data are shown as the mean ± SD (n = 4 biological replicates). Data were analyzed by two-way analysis of variance (ANOVA) with Dunnett’s multiple comparisons test (**I-N**).

Next, we analyzed the sequencing data from the lncRNA KO CRISPR screen (**Table S1**). Notably, the distribution of sgRNAs in abiraterone-treated cells on Day 28 differed significantly from that in DMSO-treated controls (**Figure 1B**). In the negative selection pool, a total of 553 lncRNAs were identified as regulators of cell growth, while 523 lncRNAs were classified as drug-resistant genes (**Figure 1C**). Conversely, in the positive selection pool, 169 lncRNAs were found to inhibit cell growth, and 717 lncRNAs were characterized as Abiraterone-sensitive genes (**Figure 1D**). Intersection analysis of the control vs. D0 and abiraterone vs. control comparisons revealed that only seven lncRNAs in the negative selection pool influenced both cell growth and abiraterone resistance (**Figure 1E**). Our study subsequently focused on lncRNAs depleted in the negative selection pool. These lncRNAs were prioritized using a multi-parameter ranking system incorporating rank score, fold change, and read count as screening criteria. The top-ranked lncRNAs demonstrated consistent reproducibility across these metrics, enhancing the robustness of our findings (**Figures 1F-1H, Figures S1D-1F**). We then selected two single-guide RNAs (sgRNAs) per gene to knock out the top three ranked lncRNAs. Functional validation confirmed that depletion of these lncRNAs significantly increased drug sensitivity in both DU145 and PC3 cells (**Figures 1I-1N, Figure S1G**).

Next, we analyzed the sequencing data from protein-coding gene KO library screen and observed significant differences in sgRNA distribution between abiraterone-treated and DMSO-treated groups (**Figure. S2A, Table S1**). The top-ranked genes, prioritized based on rank score, fold change, and read count, demonstrated consistent reproducibility across these metrics (**Figures 2A-2C**). Knockout of the top three abiraterone-sensitive genes enhanced drug sensitivity in both Du145 and PC3 cells (**Figures 2D-2I, Figure S2B**), confirming the robustness of our findings. We then performed an integrated analysis of drug-resistant lncRNAs and protein-coding genes, identifying four significantly enriched sense-antisense gene pairs (**Figure 2J**). Expression correlation analysis of these pairs revealed a positive association in prostate cancer patient samples (**Figure 2K, Figures S2C-2E**). Among these, the *BIRC6-AS1*/*BIRC6* pair emerged as a top candidate for further investigation, given that BIRC6 exhibited the highest robust rank aggregation (RRA) score and *BIRC6-AS1* showed the greatest fold change. While BIRC6 has been implicated in PCa progression and enzalutamide resistance^16^, its role in abiraterone resistance remains unexplored. To elucidate the regulatory relationship between *BIRC6-AS1* and BIRC6, we found that BIRC6 knockdown did not alter *BIRC6-AS1* expression (**Figures 2L and 2M**). Conversely, *BIRC6-AS1* depletion reduced both mRNA and protein levels of BIRC6 (**Figures 2N-2Q**).

**Figure 2.**
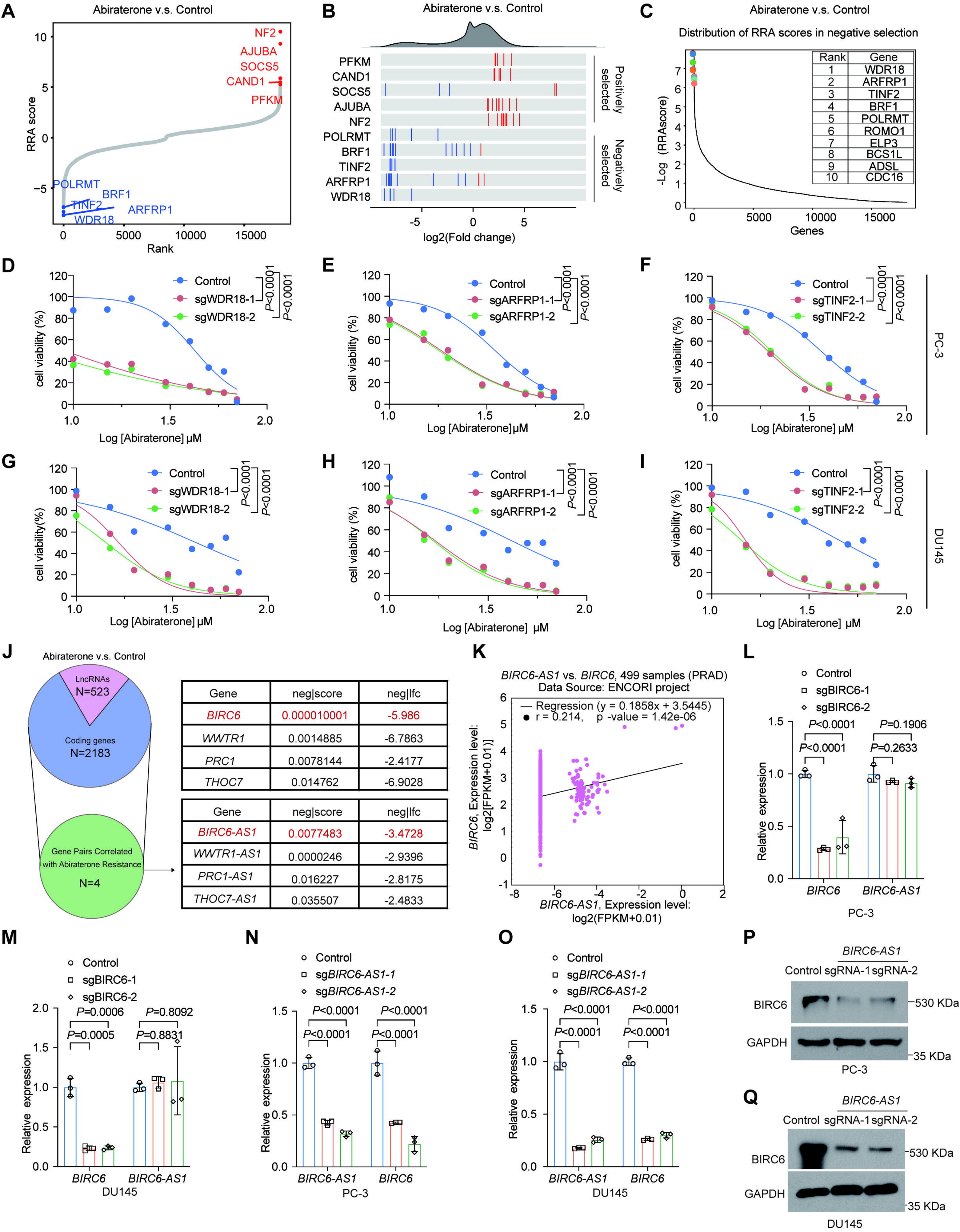
CRISPR library screening identified *BIRC6-AS1*/BIRC6 as critical paired genes for Abiraterone resistance. (**A**) MAGeCK analysis results displayed a ranking of genes based on their RRA scores from the genome-wide CRISPR Cas9 library. (**B**) Frequency distribution of log2 fold changed for all sgRNAs (top) and log2 fold change of individual sgRNAs for representative candidates (bottom). Enriched and depleted sgRNA hits were indicated by red and blue vertical bars, respectively. (**C**) The RRA score distribution plot revealed the top 10 candidate genes associated with abiraterone resistance. (**D-I**) Cell viability assays in PC3 (**D-F**) and DU145 (**G-I**) cells treated with 0-70 μM abiraterone for 48h, following transduction with either control sgRNAs or sgRNAs targeting candidate genes: *WDR18* (**D** and **G**), *ARFRP1* (**E** and **H**), and *TINF2* (**F** and **I**), Data are shown as the mean ± SD (n = 4 biological replicates). (**J**) Venn diagram showing four synergistic lncRNA-coding gene pairs (intersections) associated with abiraterone resistance, identified through CRISPR screening of lncRNA (pink) library and genome-wide (blue) library. The table quantifies screening metrics (neg|score and neg|fc) for these pairs. (**K**) Co-expression analysis of *BIRC6-AS1* and *BIRC6* in prostate samples was performed based on the ENCORI database. (**L** and **M**) qRT-PCR analysis of *BIRC6* and *BIRC6-AS1* expression in BIRC6 knockdown PC-3 (**L**) and DU145 (**M**) cells. (**N** and **O**) qRT-PCR analysis of *BIRC6-AS1* and *BIRC6* expression levels in *BIRC6-AS1* knockdown PC-3 (**N**) and DU145 (**O**) cells. (**P** and **Q**) Western blot analysis of BIRC6 protein levels in *BIRC6-AS1* knockdown PC-3 cells (**P**) and DU145 (**Q**) cells. The samples derived from the same experiment and the gels/blots were processed in parallel. Statistical differences determined by two-way ANOVA with Dunnett’s (**D-I**) or Sidak’s (**L-O**) multiple comparisons test.

Collectively, our study identified and functionally validated abiraterone-resistant lncRNAs and protein-coding genes, revealing that the sense-antisense gene pair *BIRC6-AS1*/*BIRC6* may serve as key regulators in the acquisition of abiraterone resistance in PCa cells.

### Inhibition of *BIRC6-AS1* or BIRC6 sensitized PCa cells to abiraterone treatment

To investigate the role of *BIRC6-AS1*/*BIRC6* in the therapeutic response to abiraterone in PCa cells, we first analyzed their expression profiles using multiple published RNA-seq datasets derived from PCa patient tumor samples. Our results demonstrated significant upregulation of both *BIRC6-AS1* (**Figures 3A and 3B**) and *BIRC6* (**Figures 3C and 3D**) in tumor tissues compared to paired normal controls.

**Figure 3.**
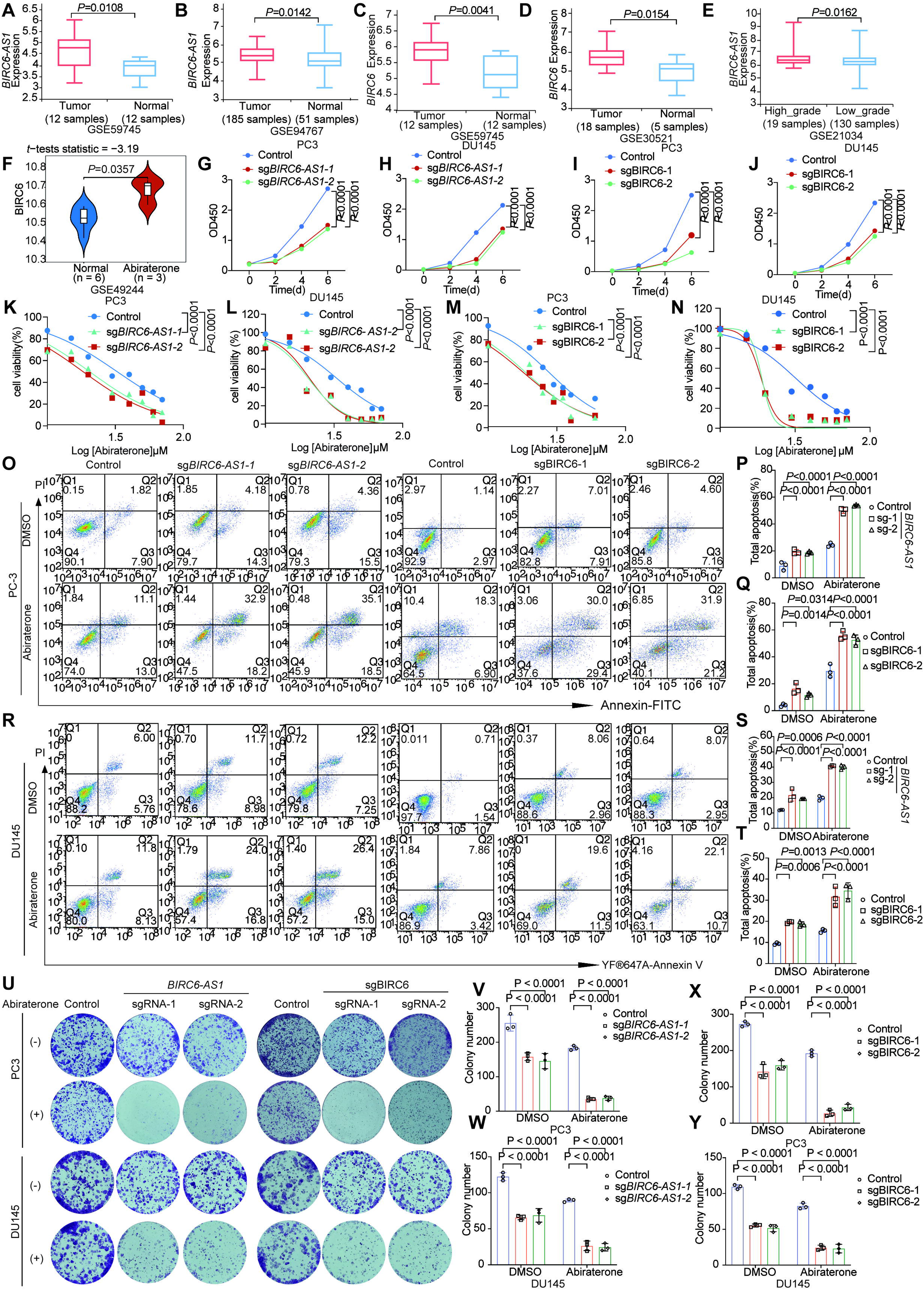
Depletion of *BIRC6-AS1* or BIRC6 potentiated abiraterone response in prostate cancer. (**A-E**) Box plots profiling *BIRC6-AS1* (**A and B, E**) and *BIRC6* (**C** and **D**) expression across prostate cancer clinical cohorts. (**A**) Primary tumors vs. normal prostate tissues (GSE59745; n=12 tumor, 12 normal). (**B**) Primary tumors vs. normal prostate tissues (GSE94767; n=185 tumor, 51 normal). (**C**) Primary tumors vs. normal prostate tissues (GSE59745; n=12 tumor, 12 normal). (**D**) Primary tumors vs. normal prostate tissues (GSE30521; n=18 tumor, 5 normal). (**E**) High-grade vs. low-grade tumors (GSE21034; n=19 high-grade, 130 low-grade). (**F**) Violin plot illustrating the expression levels of *BIRC6* in normal (n = 6) and abiraterone-treated (n = 3) prostate tissue samples from the GSE49244 dataset. (**G** and **H**) CCK-8 proliferation assays were performed in *BIRC6-AS1* knockdown PC3 (**G**) and DU145 (**H**) cells. Data are shown as the mean ± SD (n = 4 biological replicates). (**I** and **J**) CCK-8 proliferation assays were performed in BIRC6 depleted PC3 (**I**) and DU145 (**J**) cells. Data are shown as the mean ± SD (n = 4 biological replicates). (**K-N**) Cell viability assays in PC3 cells (**K** and **M**) and DU145 cells (**L** and **N**) treated with different concentrations of abiraterone (0-70 μM) for 48h, following transduction with either control sgRNAs or sgRNAs targeting *BIRC6-AS1* (**K** and **L**) and *BIRC6* (**M** and **N**). Data are shown as the mean ± SD (n = 4 biological replicates). (**O-T**) Apoptosis was analyzed by Annexin V/PI staining in PC3 (**O**) and DU145 cells (**R**) transduced with control sgRNAs or sgRNAs targeting *BIRC6-AS1* or *BIRC6*, followed by treatment with DMSO or Abiraterone (30μM) for 48h. Representative images (**O** and **R**) and quantitative statistics for *BIRC6-AS1* knockdown group (**P** and **S**) and BIRC6 depleted group (**Q** and **T**) was shown. Data are shown as the mean ± SD (n = 3 biological replicates). (**U**-**Y**) Colony formation assays in prostate cancer cells treated with DMSO or Abiraterone (30μM) for 48h. Representative images (**U**) and quantification (**V-Y**) are shown as indicated. PC3 cells (**V** and **X**) and DU145 (**W** and **Y**) cells were treated with abiraterone (30μM) for 48h, following transduction with either control sgRNAs or sgRNAs targeting *BIRC6-AS1* (**V** and **W**) and *BIRC6* (**X** and **Y**). Data were analyzed by two-way ANOVA with Dunnett’s(**G-N**) or Sidak’s (**P, Q, S, T, V-Y**) multiple comparisons test.

Notably, *BIRC6-AS1* expression was markedly higher in high-grade PCa samples than in low-grade counterparts (**Figure 3E**). Additionally, BIRC6 overexpression was observed in three abiraterone-resistant PCa samples relative to six normal samples (**Figure 3F**). Despite the limited sample size, these findings suggest a potential association between BIRC6 and abiraterone resistance. To further validate the functional contributions of *BIRC6-AS1* and *BIRC6* to drug resistance, we performed knockout experiments targeting each gene in DU145 and PC3 cell lines (**Figures S3A and 3B**). Subsequent assays evaluated effects on cellular proliferation, abiraterone sensitivity, apoptosis, and clonogenic capacity. As shown in **Figure 3G-3J**, depletion of either *BIRC6-AS1* or BIRC6 reduced cell proliferation. Consistent with our library screening results, knockout of either gene significantly augmented drug sensitivity in both PC3 and Du145 cells compared to control cells (**Figures 3K-3N**). Apoptosis and clonogenic assays further confirmed that *BIRC6-AS1* or *BIRC6* depletion increased apoptosis rates **(Figures 3O-3T**) and impaired clonogenic potential (**Figures 3U-3Y**) in abiraterone-treated Du145 or PC3 cells. Altogether, these findings corroborate our initial library screening results and establish *BIRC6-AS1*/BIRC6 as key regulators of abiraterone resistance in PCa.

### *BIRC6-AS1* stabilizes the mRNA of *BIRC6* by interacting with ILF2

Then, we investigated the regulatory mechanisms between *BIRC6-AS1* and BIRC6. We initially examined the cellular localization of *BIRC6-AS1* by employing RNA fluorescence in situ hybridization (RNA-FISH), which revealed its predominant nuclear localization in PCa cells (**Figure 4A**). Quantitative subcellular fractionation analysis confirmed that *BIRC6-AS1* is a nuclear-enriched lncRNA (**Figure 4B**). Based on the above results, we hypothesized that nuclear-localized *BIRC6-AS1* may regulate BIRC6 transcription or modulate its mRNA stability through interactions with unidentified factors. To test this hypothesis, we performed an RNA pull-down assays employing full-length *BIRC6-AS1*. A distinct band at approximately 40 kDa in the *BIRC6-AS1* sense group was excised and analyzed by mass spectrometry (**Figure 4C, Table S2**). Among the top-ranked candidates, interleukin enhancer-binding factor 2 (ILF2) was identified and validated as a *BIRC6-AS1*-interacting protein via immunoblotting (**Figure 4D and 4E**). Immunofluorescence assays further confirmed nuclear co-localization of *BIRC6-AS1* and ILF2 in PC3 and DU145 cells (**Figure 4F**), demonstrating their interaction within the nucleus.

**Figure 4.**
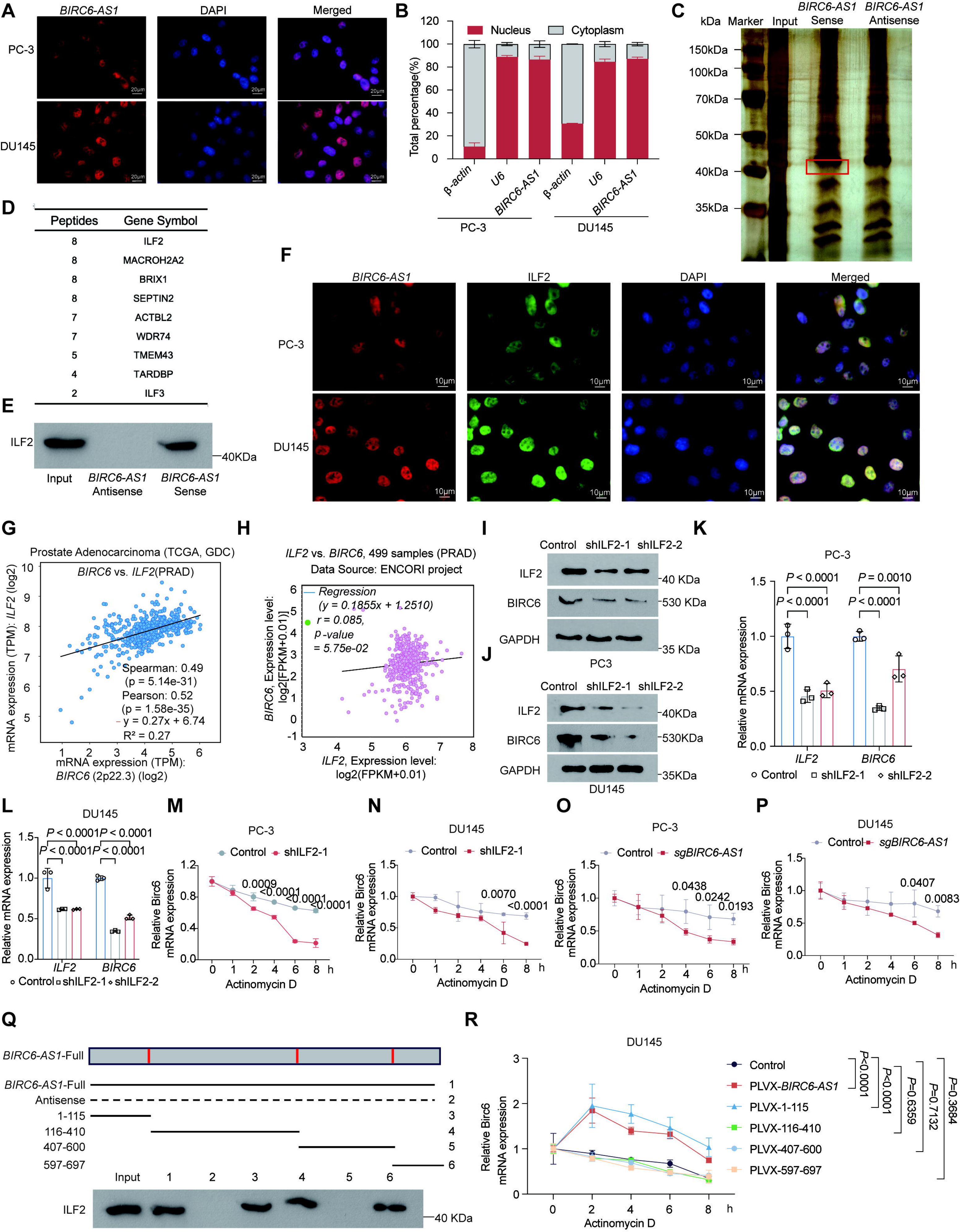
*BIRC6-AS1*-ILF2 interaction stabilized *BIRC6* mRNA. (**A**) FISH assays identified the localization of *BIRC6-AS1* in prostate cancer cells. Scale bar = 20 μm. (**B**) Subcellular localization was performed to isolate cytoplasmic and nucleoplasmic associated RNA. qRT-PCR was performed to determine the subcellular localization ratio of *BIRC6-AS1* transcripts. □*-actin* and *U6* were used as marker RNAs for quality control. (**C**) Silver-stained SDS-PAGE gel showing proteins pulled down by biotinylated *BIRC6-AS1* sense and antisense probes. The red box highlights a specific band enriched in the sense probe group. (**D**) Mass spectrometry identification of peptides recovered from the sense probe pull-down. Peptide counts and corresponding gene symbols were listed. (**E**) Western blot validation of ILF2 interaction with *BIRC6-AS1* sense probe, using input lysate and antisense probe as controls. (**F**) FISH and IF showed the co□localization of *BIRC6-AS1* and ILF2 protein in prostate cancer cells. All samples were stained with DAPI to visualize nuclei. Scale bar = 10 μm. (**G** and **H**) Co-expression analysis of *BIRC6* and *ILF2* in prostate cancer samples from TCGA (**G**) and ENCORI (**H**) databases. (**I** and **J**) Western blot analysis of BIRC6 and *ILF2* protein levels in *ILF2* depleted PC-3 (**I**) and DU145 (**J**) cells. The samples derived from the same experiment and the gels/blots were processed in parallel. (**K** and **L**) qRT-PCR analysis of *ILF2* and *BIRC6* mRNA levels in *ILF2* knockdown PC3 (**K**) and DU145 (**L**) cells. Data are shown as the mean ± SD (n = 3 biological replicates). (**M**-**P**) mRNA stability analysis was performed to assess ILF2 and *BIRC6-AS1*-mediated enhancement of *BIRC6* mRNA stability. PC3 (**M** and **O**) and DU145(**N** and **P**) cells were transduced with control shRNA/sgRNA or *ILF2*-targeting shRNAs (**M** and **N**) and *BIRC6-AS1*-targeting sgRNAs (**O** and **P**), and followed by the addition of actinomycin D (10 μg/mL) to the culture medium. Decay curves of *BIRC6* mRNA were generated using GraphPad Prism. Data are shown as the mean ± SD (n = 3 biological replicates). (**Q**) Schematic diagram (top) shows the full-length sequence of *BIRC6-AS1* and its deletion mutant variants, with nucleotide positions numerically annotated. The results (bottom) demonstrated the ILF2-*BIRC6-AS1* interaction domain, as identified by Western blot analysis. (**R**) mRNA stability analysis was performed to assess the enhancement of *BIRC6* mRNA stability mediated by *BIRC6-AS1* and its deletion mutants. DU145 cells were transduced for overexpression of *BIRC6-AS1* or its deletion mutants, and then actinomycin D (10 μg /mL) was added to the culture medium. Data are shown as the mean ± SD (n = 3 biological replicates). Data were analyzed by two-way ANOVA with Dunnett’s (**K, L, R**) or Sidak’s (**M-P**) multiple comparisons test.

Subsequently, we evaluated the expression correlation between *ILF2* and *BIRC6* in TCGA and ENCORI prostate tumor cohorts, revealing a significant positive correlation (**Figures 4G and 4H**). Similarly, in the GSE85672 cohort of 30 abiraterone-treated CRPC tumors, BIRC6 expression positively correlated with ILF2 instead of ILF3 (**Figure S4A**). To further validate the regulatory interplay between ILF2 and BIRC6, we reduced the expression of ILF2 in both DU145 and PC3 cells, which reduced BIRC6 expression at both the protein and mRNA levels (**Figures 4I-4L**). Previous studies have reported that ILF2, in complex with ILF3, regulates nuclear mRNA distribution, DNA repair, and mRNA stability^17,18^. Consistent with these findings, ILF2 depletion in PC3 and DU145 cells accelerated *BIRC6* mRNA degradation following actinomycin D treatment compared to controls (**Figures 4M and 4N**). A similar acceleration in *BIRC6* mRNA decay was observed upon *BIRC6-AS1* knockout (**Figures 4O and 4P**). To identify the *BIRC6-AS1* region responsible for ILF2 binding, we generated truncated variants and performed binding assays. The findings revealed that two fragments (1-410 nt and 597-697 nt) of *BIRC6-AS1* were capable of sufficient binding to ILF2 (**Figure 4Q**). Furthermore, we investigated the effects of the truncated *BIRC6-AS1* variants on the stability of *BIRC6* mRNA in DU145 cells following treatment with actinomycin D. The results revealed that overexpression of full-length *BIRC6-AS1* or 1-115 nt fragment enhanced the stability of *BIRC6* mRNA compared to the control cells (**Figure 4R**).

Collectively, these findings demonstrated that *BIRC6-AS1* modulates the stabilization of *BIRC6* mRNA by interacting with ILF2.

### Depletion of *BIRC6-AS1* or BIRC6 potentiated abiraterone-induced DNA damage in PCa cells

To investigate the downstream pathways regulated by *BIRC6-AS1*/BIRC6 in abiraterone-treated cells, we performed Kyoto Encyclopedia of Genes and Genomes (KEGG) pathway analysis utilizing a panel of 523 previously identified abiraterone-resistant genes. We observed an enrichment of genes within the DNA replication and nucleotide excision repair pathways (**Figure 5A**), indicating that abiraterone treatment may induce DNA damage in PCa cells. Consequently, we employed the DR-GFP and EJ5-GFP reporter systems to quantify DNA repair activity. An augmentation in GFP fluorescence intensity served as an indicator that abiraterone treatment facilitated both non-homologous end joining (NHEJ) and homologous recombination (HR) repair processes in PC3 and DU145 cells (**Figure 5B-5D, Figure S5A and 5B**). Next, we assessed whether *BIRC6-AS1* or BIRC6 knockout exacerbates abiraterone-induced DNA damage in PCa cells. Using alkaline comet assays, we observed that *BIRC6-AS1* or *BIRC6* knockout cells exhibited longer comet tails compared to controls, with a more pronounced effect under abiraterone treatment (**Figures 5E-5G**). Similarly, γH2AX foci formation—a marker of DNA double-strand breaks (DSBs) —was significantly elevated in *BIRC6-AS1* or *BIRC6*-deficient PCa cells, particularly when combined with abiraterone (**Figures 5H-5J**).

**Figure 5.**
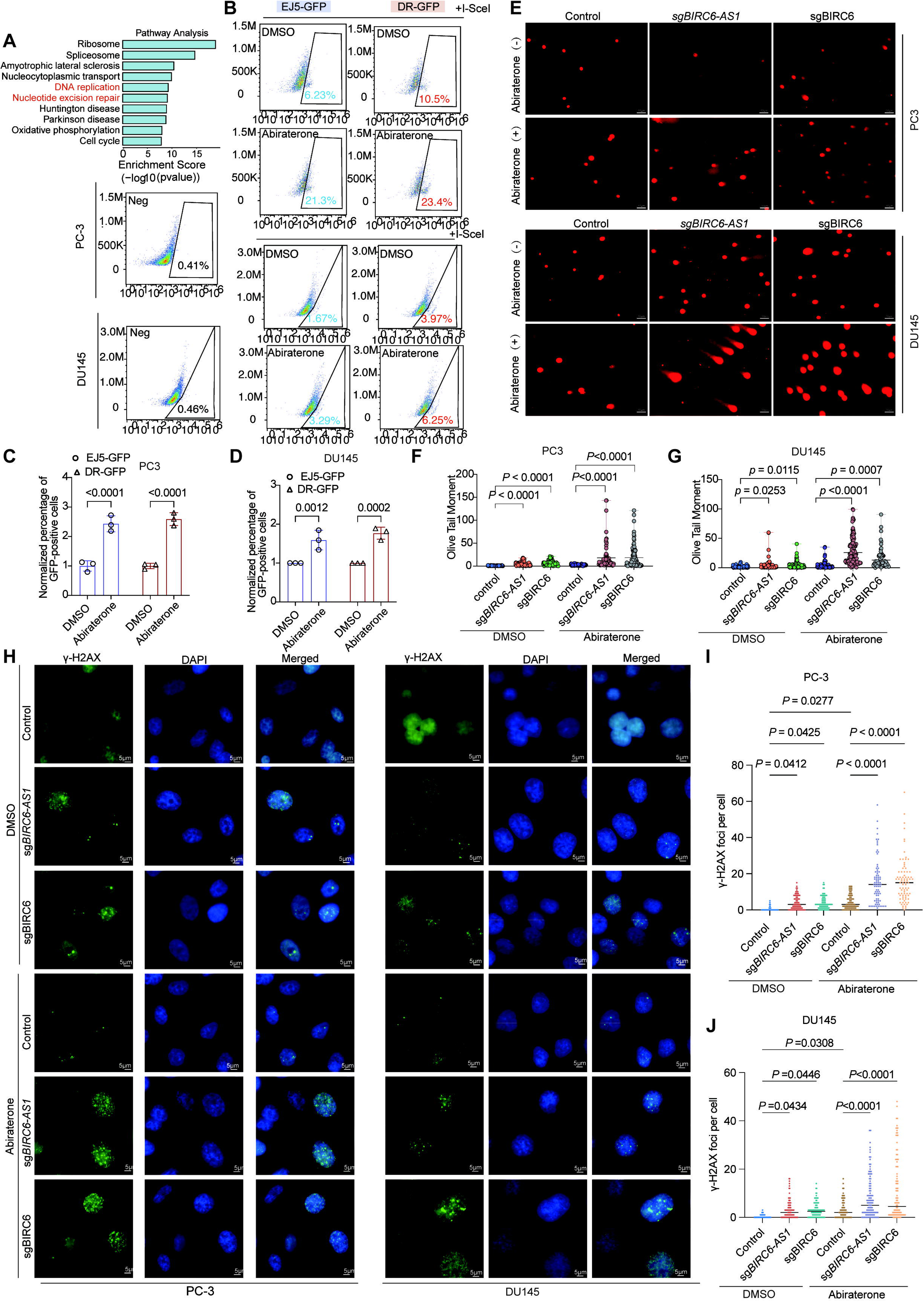
*BIRC6-AS1*/BIRC6 inhibition enhances abiraterone-triggered DNA damage response in PCa cells. (**A**) KEGG pathway analysis of abiraterone resistance-associated genes in PC3 cells (identified via genome-wide CRISPR screening). (**B-D**) Assessment of non-homologous end joining (NHEJ) and homologous recombination (HR) repair efficiency in PC3 and DU145 cells cultured with or without abiraterone (30 μM). Representative images (**B**) and quantification (**C** and **D**) were shown as indicated. (**E-G**) Representative neutral comet assay images of PC3 and DU145 cells with *BIRC6-AS1* or *BIRC6* depleted, in the presence of abiraterone (30 μM). Representative images (**E**) and quantification (**F** and **G**) were shown as indicated. Data from 90 cells were averaged to calculate tail moment. Scale bar = 50 μm. **(H-J**) Immunofluorescence staining of γ-H2AX in *BIRC6-AS1*/BIRC6 knockdown PC3 (**H** and **I**) cells and DU145 (**H** and **J**) cells cultured with or without abiraterone (30 μM). Representative images (**H**) and quantification (**I** and **J**) were shown as indicated. Data from 100 cells were averaged to calculate. Scale bar = 5 μm. Data were analyzed by two-way ANOVA (**C** and **D**) with Sidak’s multiple comparisons test or one-way ANOVA (**F, G, I, J**) with Tukey’s multiple comparisons test.

To validate these findings *in vivo*, we established xenograft models by subcutaneously inoculating nude mice with PC3 cells harboring *BIRC6-AS1* knockout, BIRC6 knockout (sgBIRC6), or control cells (1×10^6^ cells per group) The schematic outline of the experiment is depicted in **Figure 6A**. The mice were administered abiraterone (150 mg/kg) via oral gavage every other day until study termination. Notably, abiraterone treatment synergized with *BIRC6-AS1* or *BIRC6* knockout to suppress tumor growth (**Figures 6B-6D**). Immunohistochemical (IHC) analysis of xenograft tumors confirmed reduced BIRC6 protein expression in *BIRC6-AS1* and BIRC6 knockdown groups, validating the regulatory role of *BIRC6-AS1* in modulating BIRC6 levels (**Figures 6E and 6F**). Moreover, γH2AX levels were elevated in tumors with *BIRC6-AS1* or BIRC6 depletion, and abiraterone treatment further exacerbated γH2AX levels (**Figures 6E and 6G**).

**Figure 6.**
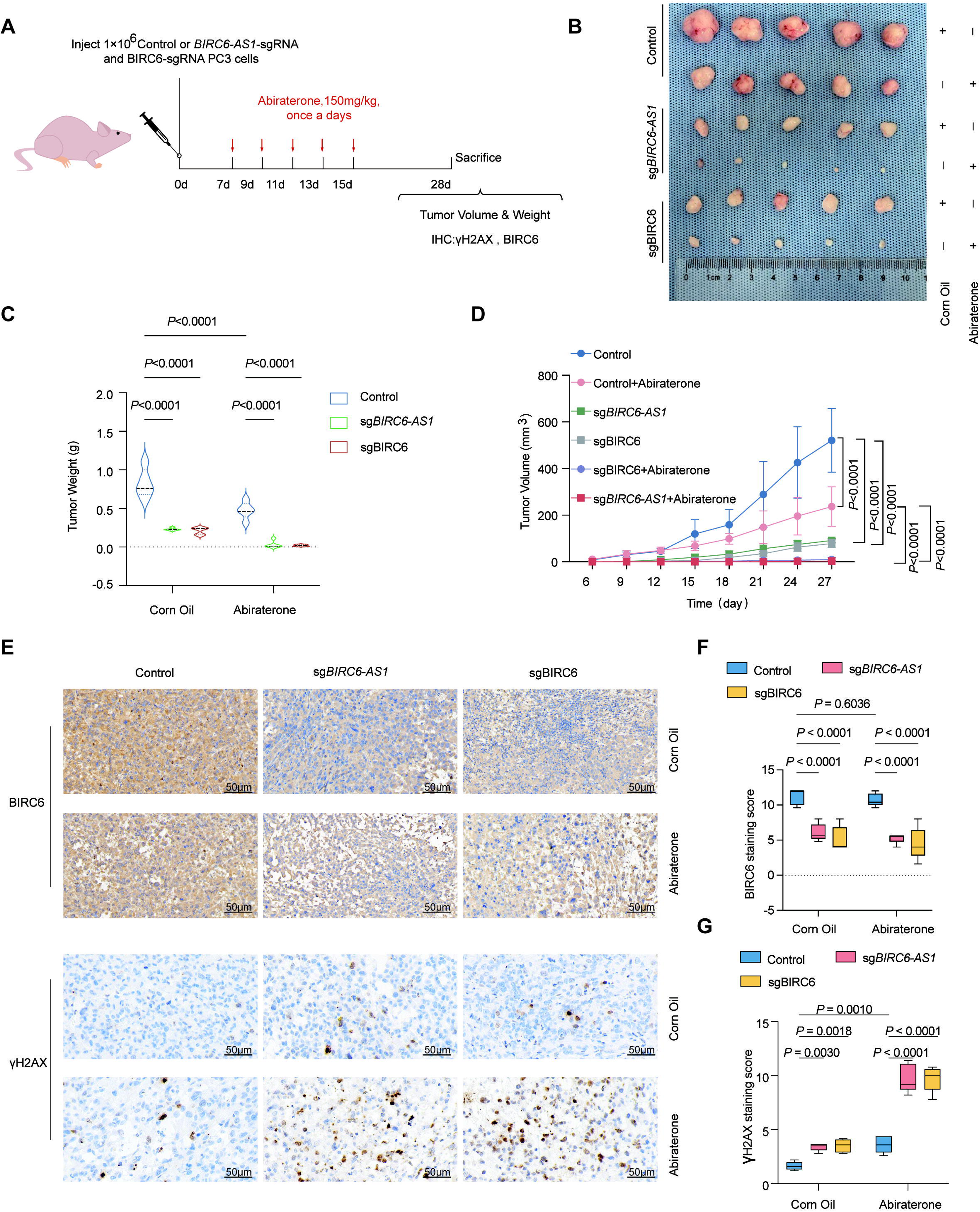
Depletion of *BIRC6-AS1*/BIRC6 increased abiraterone-induced γH2AX foci formation in vivo. (**A**) Schematic diagram of the in vivo experimental workflow. 1×10□ PC3 sg*BIRC6-AS1* cells, sgBIRC6 cells, and control cells were subcutaneously injected into nude mice, respectively. Mice received abiraterone (150 mg/kg) via gavage every other day during the experiment, and tumor tissues were harvested on day 28 post-inoculation. (**B** and **C**) Representative image (**B**) and tumor weight data (**C**) for each indicated group at the experimental endpoint. Values are presented as mean ± SD (n = 5 tumors). (**D**) Tumor growth curves. Data are presented as mean ± SD (n = 5 tumors). (**E**-**G**) Immunohistochemical (IHC) staining of BIRC6 (**E** and **F**) and γH2AX (**E** and **G**) was performed in isolated tumor tissues. Data are presented as mean ± SD (n = 5 tumors). Each tumor section was randomly selected for 5 magnification fields. Scale bars = 50 μm. Statistical differences were determined using two-way ANOVA with Sidak’s multiple (**C, F, G**) or Tukey’s (**D**) comparisons test.

Taken together, these findings indicated that abiraterone treatment induces DNA damage in PCa cells, and knockout of either *BIRC6-AS1* or BIRC6 further exacerbates this DNA-damaging effect.

### BIRC6 interacted with A20 and facilitated its ubiquitination and degradation in prostate cancer

To elucidate the downstream molecular mechanisms by which *BIRC6-AS1*/BIRC6 modulate abiraterone resistance, we examined which DNA repair pathway was regulated by these two genes. The results demonstrated that decreasing the expression of either *BIRC6-AS1* or BIRC6 predominantly suppressed the NHEJ repair activity other than HR repair process (**Figure 7A, Figure S6A**). Given that the BIRC6 functions as an E2/E3 ubiquitin ligase, we hypothesized that BIRC6 may probably regulate the NHEJ repair pathway through ubiquitination of key factors involved in this process. Then, using the UbiBrowser2.0, we predicted BIRC6-interacting proteins and identified three potential candidates, including OTUB1, OTUB2 and A20 (**Figure 7B**). Among these, A20 has been previously reported to suppress NHEJ activity and is essential for the disassembly of 53BP1 at DNA damage foci ^19^. Knockout of either *BIRC6-AS1* or BIRC6 enhanced the nuclear localization of A20, and this effect was further potentiated in PCa cells treated with abiraterone (**Figures 7C and 7D**). Immunofluorescence microscopy revealed that knockout of *BIRC6-AS1* or BIRC6 promoted 53BP1 disassembly from DNA damage foci (**Figures 7E and 7F**). These results suggested that BIRC6 probably regulates the NHEJ repair pathway via A20, prompting us to further investigate the regulatory interplay between BIRC6 and A20.

**Figure 7.**
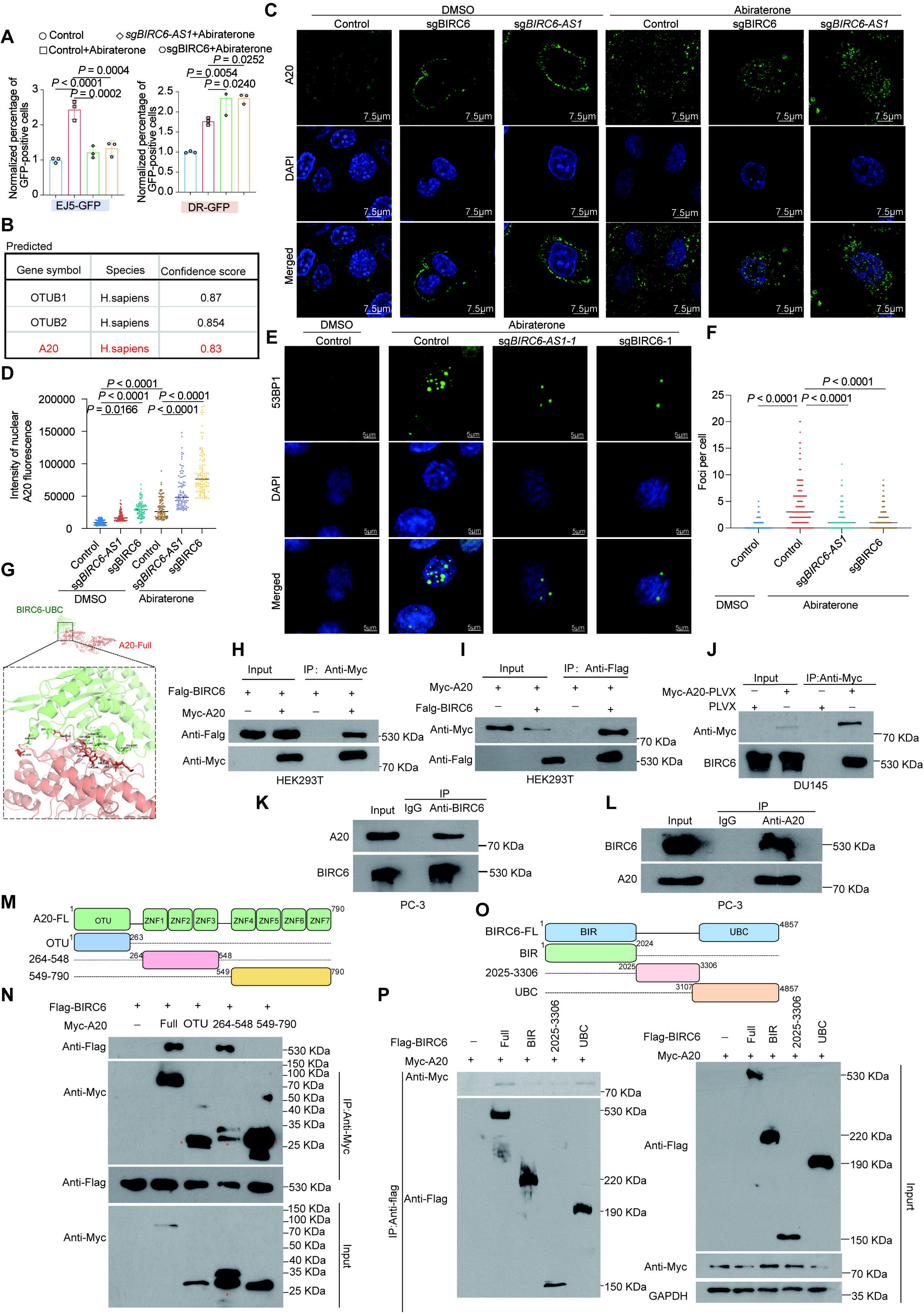
BIRC6 physiologically interacted with A20. (**A**) Assessment of non -homologous end joining (NHEJ) and homologous recombination (HR) repair efficiency in *BIRC6-AS1*/BIRC6 knockdown PC3 and DU145 cells cultured with or without abiraterone (30 μM). (**B**) Table showing the top 3 predicted protein interactors of BIRC6. **C-D:** Immunofluorescence staining of A20 in *BIRC6-AS1* or BIRC6 depletion DU145 cells cultured with or without abiraterone (30 μM). Representative images (**C**) and quantification (**D**) were shown as indicated. Data from 100 cells were averaged to calculate. The nucleus was stained with DAPI. Scale bar = 7.5 μm. (**E** and **F**) Immunofluorescence staining of 53BP1 in *BIRC6-AS1* or BIRC6 knockout DU145 cells treated with or without abiraterone (30 μM). Representative images (**E**) and quantification (**F**) are shown as indicated. Data from 100 cells were averaged to calculate. The nucleus was stained with DAPI. Scale bar = 5 μm. (**G**) Protein-protein docking model between A20 (red) and BIRC6-UBC (green). Detailed view shows the interaction interface with rediction as to the most probable amino acid of interaction (A20 protein: K337). (**H** and **I**) HEK293T cells were transfected with expression plasmids for Myc-A20 and Flag-BIRC6 as indicated. Whole-cell lysates were extracted and immunoprecipitated with anti-Myc (**H**) or anti-Flag (**I**) antibodies. The samples derived from the same experiment and the gels/blots were processed in parallel. (**J**) Myc-A20-pLVX-overexpressing DU145 cells or control cells were immunoprecipitated with anti-Myc antibody and then subjected to western blot analysis. The samples derived from the same experiment and the gels/blots were processed in parallel. (**K** and **L**) Whole-cell lysates of PC3 cells were collected, and immunoprecipitations were performed with anti-BIRC6 (**K**) or Anti-A20 (**L**) antibodies, followed by western blot with indicated antibodies. The samples derived from the same experiment and the gels/blots were processed in parallel. (**M** and **O**) Schematic diagram of full-length A20 (**M**) and BIRC6 (**O**) and their respective truncated mutants was shown. (**N**) HEK293T cells were transfected with full-length A20 or its truncated mutants, as well as full-length BIRC6. Lysates were prepared and subjected to immunoprecipitation with anti-Myc antibodies to map the binding region of BIRC6 with A20. The samples derived from the same experiment and the gels/blots were processed in parallel. (**P**) HEK293T cells were transfected with full-length BIRC6 or its truncated mutants, as well as full-length A20. Lysates were prepared and subjected to immunoprecipitation with anti-Flag antibodies. The samples derived from the same experiment and the gels/blots were processed in parallel. Data were analyzed by one-way ANOVA with Dunnett’s multiple(**A**) or Tukey’s (**D** and **F**) comparisons test.

To characterize the the structural basis of the BIRC6-A20 interaction, molecular docking was performed. As shown in the model (**Figure 7G**), the UBC domain of BIRC6 formed a spatial binding complex with full-length A20. To confirm the interaction between BIRC6 and A20, we co-transfected Flag-BIRC6 and Myc-A20 into HEK293T cells and performed Co-IP assay using anti-Myc or anti-Flag antibody. The results showed that BIRC6 interacts with A20 (**Figures 7H and 7I**). Semi-endogenous and endogenous Co-IP assay further verified that BIRC6 physiologically interacts with A20 (**Figures 7J-7L**). To determine which region of A20 is required for BIRC6 binding, Myc-tagged full-length A20 and deletion mutants were co-transfected into HEK293T cells with BIRC6 expression vector. Immunoprecipitations were performed using anti-Myc antibody. The results indicated that the 264-548 aa functional domain of A20 was essential for the binding of BIRC6 (**Figures 7M and 7N**). We also verified which region of BIRC6 can bind to A20, and found that the UBC domain of BIRC6 was essential for A20 binding (**Figures 7O and 7P**).

To explore the relationship between *BIRC6-AS1*/BIRC6 and A20, we first investigated whether *BIRC6-AS1*/BIRC6 affected expression levels of A20. The results demonstrated that *BIRC6-AS1* or BIRC6 knockdown unchanged mRNA levels of *A20*, but increased protein levels of A20 (**Figures 8A and 7B, Figures S8A and 8B**). Next, we found that the expression level of A20 was gradually decreased when the amount of BIRC6 increased in HEK293T cells (**Figure 8C**). To investigate the molecular mechanism by which *BIRC6-AS1*/BIRC6 regulated A20 protein level, we carried out the Cycloheximide (CHX) chase assays. The results showed that knockdown of *BIRC6-AS1* or BIRC6 prolonged the half-life of endogenous A20 protein in DU145 cells (**Figures 8D and 8E**). In contrast, BIRC6 overexpression accelerated the protein degradation of A20 and shortened its half-life (**Figure 8F**). Further domain-mapping experiments demonstrated that BIRC6 mutants lacking the A20 binding domain failed to shorten the half-life of A20 protein, whereas the isolated UBC domain of BIRC6 (BIRC6-UBC) could weaken A20 protein stability **(Figures 8G-8I**). These findings indicated that BIRC6 promoted the protein degradation of A20 depending on the interaction of BIRC6 with A20.

**Figure 8.**
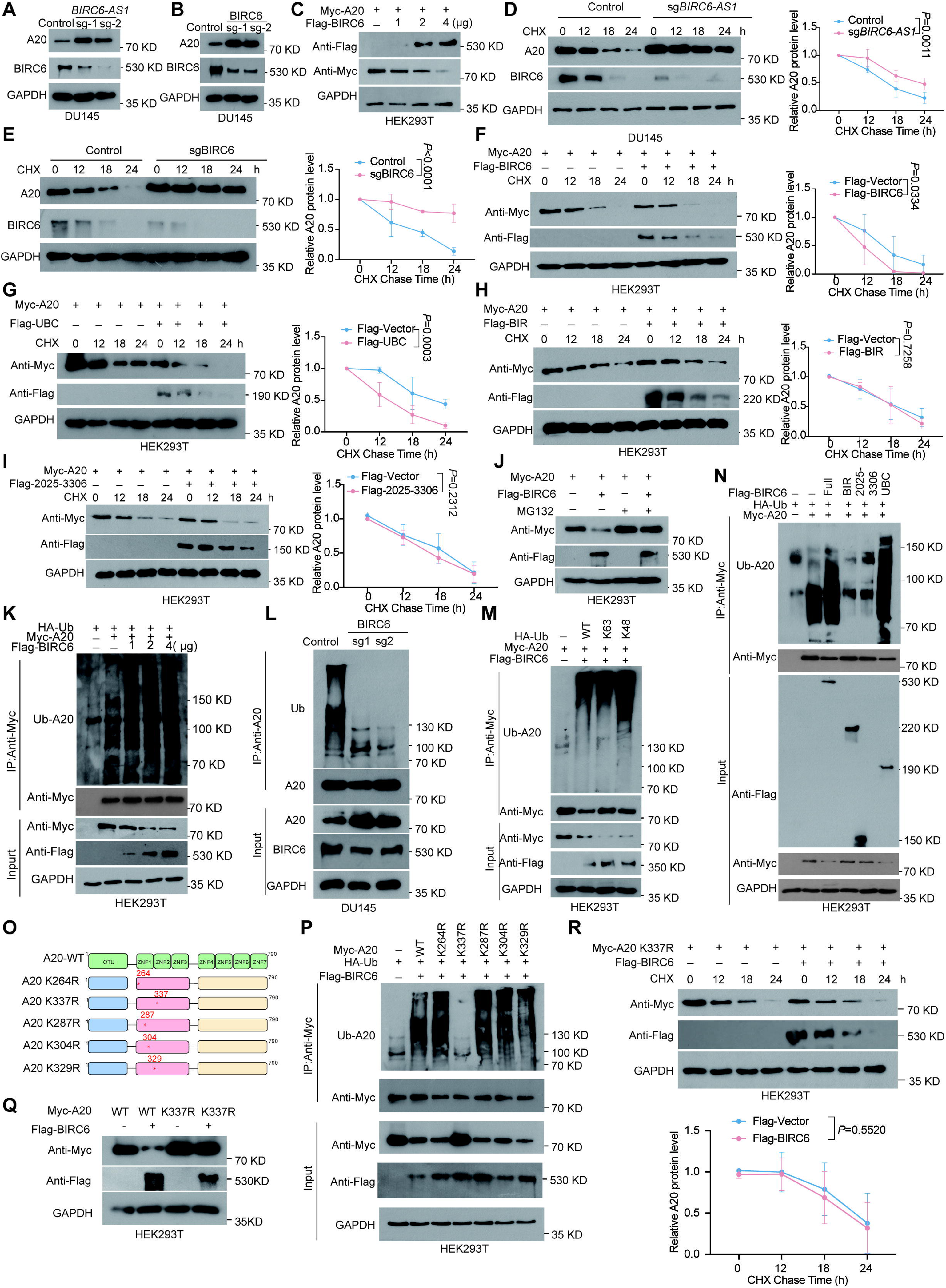
BIRC6 facilitated the K48-linked ubiquitination and degradation of A20 at the K337 residue. (**A** and **B**) Western blot analysis of A20 and BIRC6 protein levels in DU145 cells with *BIRC6-AS1* (**A**) or BIRC6 knockout (**B**). The samples derived from the same experiment and the gels/blots were processed in parallel. (**C**) HEK293T cells were co-transfected with Myc-A20 and increasing amounts of Flag-BIRC6 expression plasmids as indicated, and A20 protein level was analyzed by Western blot. GAPDH was used as a loading control. The samples derived from the same experiment and the gels/blots were processed in parallel. (**D** and **E**) DU145 cells with *BIRC6-AS1* (**D**) or BIRC6 (**E**) depletion were treated with 50 μg /mL cycloheximide (CHX) for the indicated times, and A20 protein stability was assessed by Western blot. Data are presented as mean ± SD from three independent experiments. The samples derived from the same experiment and the gels/blots were processed in parallel. (**F-I**) HEK293T cells co-transfected with full-length A20 and either full-length BIRC6 or its truncated forms were treated with 50 μg/mL CHX for the indicated times. Western blotting evaluated A20 stability mediated by full-length BIRC6 (**F**) or mutants UBC (**G**), BIR (**H**) and 2025-3306(**I**). Data are presented as mean ± SD from three independent experiments. The samples derived from the same experiment and the gels/blots were processed in parallel. (**J**) HEK293T cells co-transfected with Myc-A20 and Flag-BIRC6 expression plasmids were treated with or without MG132 (10 μM) for 8 hours. Cell lysates were subjected to immunoblot with anti-Flag or anti-Myc antibody. The samples derived from the same experiment and the gels/blots were processed in parallel. (**K**) HA-Ub, Myc-A20 and various doses of Flag-BIRC6 expression plasmids were co-transfected into HEK293T cells. Immunoprecipitation was performed with anti-Myc antibody, and subjected to western blot with anti-HA, Myc or Flag antibody. The samples derived from the same experiment and the gels/blots were processed in parallel. (**L**) The whole cell lysates were prepared from BIRC6 depletion DU145 cells or control cells, and immunoprecipitations were performed with anti-A20 antibodies, followed by western blot with indicated antibodies. The samples derived from the same experiment and the gels/blots were processed in parallel. (**M**) HEK293T cells were transfected with Myc-A20, Flag-BIRC6 together with HA-Ub Wild-Type and its mutants plasmids (K63, K48). Immunoprecipitation was performed with anti-Myc antibody, and subjected to western blot with anti-HA, Myc or Flag antibody. The samples derived from the same experiment and the gels/blots were processed in parallel. (**N**) HEK293T cells were transfected with HA-Ub and Flag-BIRC6 or truncated plasmids together with Myc-A20 or control vectors. Immunoprecipitation was performed with anti-Myc antibody, and subjected to western blot with anti-HA, Myc or Flag antibody. The samples derived from the same experiment and the gels/blots were processed in parallel. (**O**) Schematic diagram of full-length A20 and its mutants was shown. (**P**) HEK293T cells were transfected with HA-Ub, Flag-BIRC6 together with Myc-A20 Wild-Type and its mutants plasmids as indicated. Immunoprecipitation was performed with anti-Myc antibody, and subjected to western blot with anti-HA, Myc or Flag antibody. The samples derived from the same experiment and the gels/blots were processed in parallel. (**Q**) HEK293T cells were co-transfected with Myc-A20 Wild-Type or mutant Myc-A20 K337R together with Flag-BIRC6 or control vectors, followed by Western blot analysis of A20 protein level. The samples derived from the same experiment and the gels/blots were processed in parallel. (**R**) HEK293T cells transfected with Myc-A20 K337R and full-length BIRC6 or control vectors were treated with 50 μg/mL CHX for the indicated times. Western blot analysis assessed BIRC6-mediated stabilization of the A20 mutant (A20 K337R). The samples derived from the same experiment and the gels/blots were processed in parallel. Data were analyzed by Ordinary two-way ANOVA (**E-I, R**).

We next determined whether BIRC6-mediated A20 degradation occurs via the ubiquitin-proteasome system (UPS). Treatment with the proteasome inhibitor MG132 significantly attenuated BIRC6-induced A20 degradation (**Figure 8J**), suggesting that BIRC6 attenuated A20 protein stability through UPS-dependent proteolysis. To further confirm the results, we co-expressed HA-ubiquitin, Myc-A20 or Flag-BIRC6 in HEK293T cells. A20 protein was immunoprecipitated by Myc antibody, and the ubiquitinated protein was detected with HA antibody. The results indicated that BIRC6 overexpression could strongly strengthen the ubiquitination of A20 protein in a dose-dependent manner (**Figure 8K**). Accordingly, BIRC6 knockdown suppressed the ubiquitination of A20 protein in DU145 cells (**Figure 8L**). To characterize the type of polyubiquitin linkages involved, we co-transfected A20 with ubiquitin mutants K48- or K63-linked chains (K48-Ub or K63-Ub). Immunoprecipitate and immunoblotting demonstrated that BIRC6 promoted both K48- and K63-linked polyubiquitination of A20 (**Figure 8M**). Domain-mapping experiments further revealed that the UBC domain of BIRC6 (BIRC6-UBC) was both necessary and sufficient to enhance A20 ubiquitination, whereas deletion mutants lacking the UBC domain (BIR or 2025-3306) failed to induce A20 polyubiquitination (**Figure 8N**). These data indicated that BIRC6 induces A20 degradation and ubiquitination through its UBC domain. Guided by molecular docking predictions of key BIRC6-UBC-A20 interface residues, we identified five candidate lysine residues on A20 as potential ubiquitination sites (**Figure 8O**). Site-directed mutagenesis of these residues revealed that the K337R mutation specifically abolished BIRC6-mediated A20 ubiquitination (**Figure 8P**). Moreover, the K337R mutation markedly abrogated the inhibitory effect of BIRC6 on A20, and unchanged the degradation rate of A20 protein (**Figures 8Q and 8R**). Taken together, these findings indicated that BIRC6 facilitates the K48-linked ubiquitination and proteasomal degradation of A20 at the K337 residue.

### A20 knockdown reversed abiraterone sensitivity caused by *BIRC6-AS1* depletion, both in vitro and in vivo

To validate whether A20 functions as a key downstream factor in the *BIRC6-AS1*/BIRC6 axis mediating abiraterone resistance, we reduced A20 expression in *BIRC6-AS1* knockout DU145 cells (**Figure S8A**). The proliferation assays revealed that knockdown of A20 augmented the proliferative capacity of DU145 cells (**Figure 9A, Figure S8B**), effectively rescuing the decline in proliferation observed in *BIRC6-AS1* knockout cells (**Figure 9B**). Subsequently, we assessed whether A20 could revert the drug sensitivity phenotype of *BIRC6-AS1* knockout cells by evaluating the IC50 values and clonogenic potential of cells across distinct experimental groups. Specifically, A20 knockdown elevated the IC50 value and clonogenic ability of DU145 cells in the presence of abiraterone, thereby counteracting the heightened drug sensitivity induced by *BIRC6-AS1* knockout (**Figures 9C-9F**). Furthermore, we utilized immunofluorescence experiments to quantify 53BP1 accumulation at DNA damage foci in distinct cell populations. Our findings indicated that *BIRC6-AS1* knockout diminished 53BP1 recruitment, whereas A20 knockdown potentiated 53BP1 accumulation at DNA damage foci. Notably, A20 depletion further enhanced 53BP1 recruitment in *BIRC6-AS1* knockout cells, consistent with our previous observations (**Figures 9G and 9H**).

**Figure 9.**
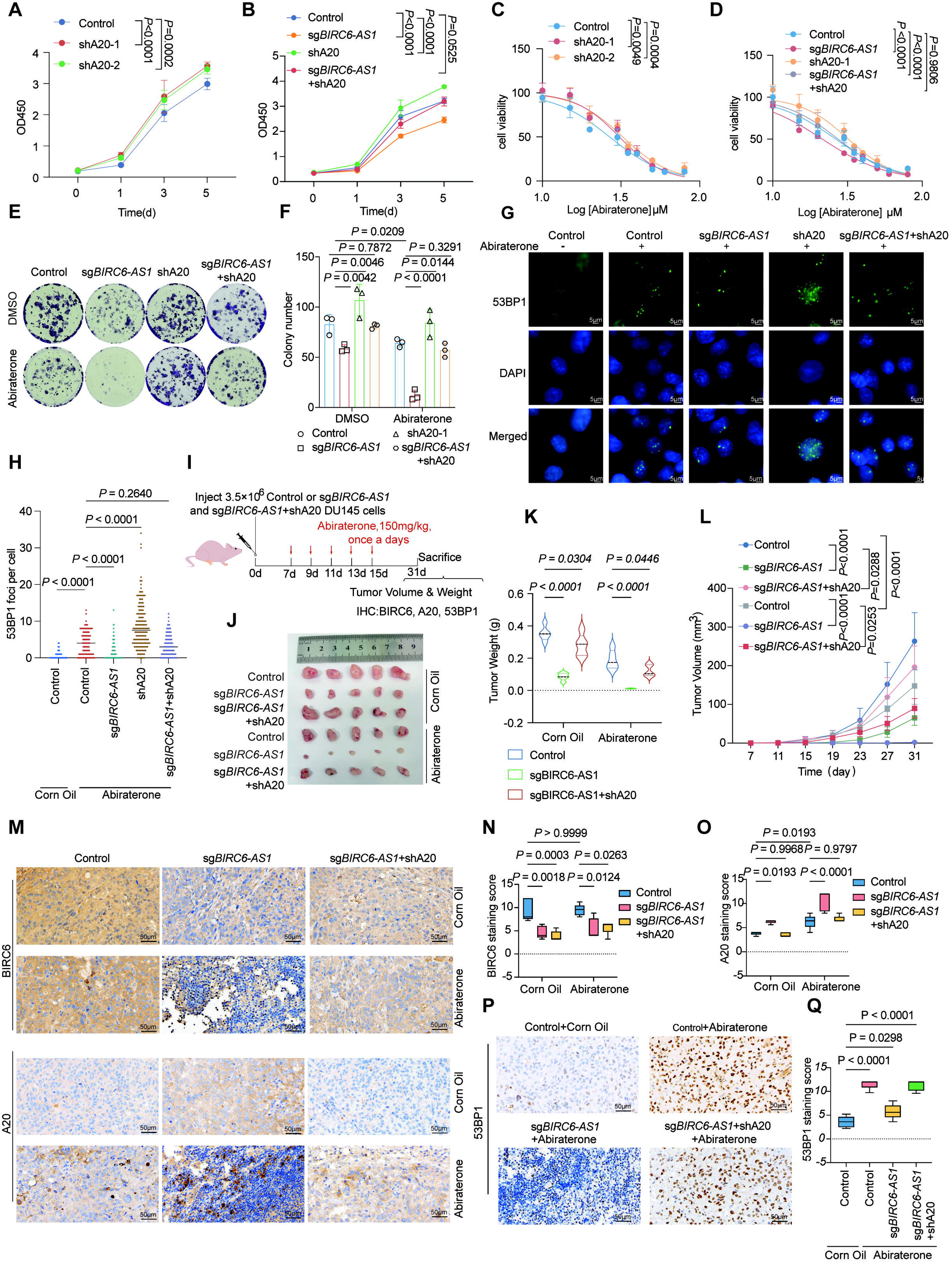
A20 silencing abrogated the abiraterone sensitization effect induced by *BIRC6-AS1* depletion both in vitro and in vivo. (**A**) The cell proliferation was measured in A20 knockdown DU145 cells by CCK8 assay. Data are shown as the mean ± SD (n = 4 biological replicates). (**B**) Cell viability of DU145 cells (Control, sg *BIRC6-AS1*, shA20, and sg*BIRC6-AS1*+shA20 groups) was evaluated by CCK-8 assay for the indicated times. Data are shown as the mean ± SD (n = 4 biological replicates). (**C**) Cell viability of A20 suppressed DU145 cells or control cells was assessed by CCK-8 in the presence of varying concentrations of abiraterone (0–70μM) for 48h. Data represent mean ± SD (n=4 biological replicates). (**D**) DU145 cells (Control, sg*BIRC6-AS1*, shA20, and sg*BIRC6-AS1*+shA20 groups) were treated with increasing concentrations of abiraterone (0–70μM) for 48h, then the cell viability were analyzed by CCK8 assay. Data represent mean ± SD (n=4 biological replicates). (**E** and **F**) DU145 cells (Control, sg*BIRC6-AS1*, shA20, and sg*BIRC6-AS1*+shA20 groups) were treated with or without abiraterone (30 μM) for 48h. colony formation ability (n =4 biological replicates) were analyzed by Crystal Violet Aqueous Solution staining. Representative images (**E**) and quantification (**F**) were shown as indicated. (**G** and **H**) Immunofluorescence staining of 53BP1 in DU145 cells (Control, sgBIRC6, shA20, and sgBIRC6+shA20 groups) cultured with or without abiraterone (30 μM) for 48h. Representative images (**G**) and quantification (**H**) were shown as indicated. Data from 100 cells were averaged to calculate. The nucleus was stained with DAPI. Scale bar = 5 μm. (**I**) Schematic diagram of the in vivo experimental workflow. 3.5×10□ sg*BIRC6-AS1* DU145 cells, sg*BIRC6-AS1*+shA20 DU145 cells, and control cells were subcutaneously injected into nude mice (n=5 per group), respectively. Mice received abiraterone (150 mg/kg) via gavage every other day during the experiment, and tumor tissues were harvested on day 31 post-inoculation. (**J** and **K**) Representative image (**J**) and tumor weight data (**K**) for each indicated group at the experimental endpoint. Values are presented as mean ± SD (n = 5 tumors). (**L**) Tumor growth curves. Data are presented as mean ± SD (n = 5 tumors). (**M-Q**) Immunohistochemical (IHC) staining of BIRC6 (**M** and **N**), A20 (**M** and **O**) and 53BP1 (**P** and **Q**) was performed in isolated tumor tissues. Data are presented as mean ± SD (n = 5 tumors). Each tumor section was randomly selected for 5 magnification fields. Scale bars = 50 μm. Data were analyzed by two-way ANOVA with Dunnett’s multiple (**A-D, K**) or Tukey’s (**F, L, N, O**) comparisons test and one-way ANOVA with Dunnett’s multiple (**H**) or Tukey’s (**Q**) comparisons test.

To evaluate the in vivo relevance of the *BIRC6-AS1*/A20 axis in abiraterone resistance, we established subcutaneous xenograft tumor models by inoculating 3×10^6^ DU145 cells into nude mice. Experimental groups included *BIRC6-AS1* knockout, dual *BIRC6-AS1*/A20-deficient, and control cells. Each group was distinctly inoculated, and the experimental design is schematically depicted in **Figure 9I**. The mice were administered abiraterone (150 mg/kg) via oral gavage every other day throughout the study duration. abiraterone effectively suppressed tumor growth in xenografts derived from *BIRC6-AS1* knockout DU145 cells. However, concurrent knockdown of A20 in *BIRC6-AS1* knockout xenografts attenuated the enhanced drug sensitivity initially induced by *BIRC6-AS1* depletion (**Figures 9J-9L**). Immunohistochemical (IHC) staining was employed to quantify the protein levels of BIRC6 and A20, as well as the nuclear accumulation of 53BP1, in the xenograft tumor tissues. The findings demonstrated that xenograft tumors derived from *BIRC6-AS1* knockout cells displayed reduced BIRC6 protein level (**Figures 9M and 9N**), elevated A20 expression levels (**Figures 9M and 9O**), and decreased nuclear accumulation of 53BP1. Consistent with in vitro findings, xenograft tumors with concurrent downregulation of both *BIRC6-AS1* and A20 revealed that A20 suppression reversed the reduced nuclear accumulation of 53BP1 caused by *BIRC6-AS1* knockout (**Figures 9P and 9Q**).

In summary, A20 emerges as a crucial downstream mediator of the *BIRC6-AS1*/BIRC6 regulatory axis, modulating abiraterone resistance in PCa cells through regulation of the NHEJ DNA repair pathway.

## Discussion

In the present study, by using genome-wide CRISPR/Cas9 library screening system, we identified 523 lncRNAs and 2183 protein-coding genes association with abiraterone-resistance. By integrated analysis, we found that *BIRC6-AS1*/*BIRC6* might be the crucial paired sense-antisense genes promoting abiraterone resistance. KEGG analysis of these resistant genes revealed a high enrichment in DNA repair pathways, suggesting the close association between the development of abiraterone resistance and DNA damage repair. Subsequently, we conducted an in-depth study on the relationship between *BIRC6-AS1*/BIRC6 and abiraterone resistance, demonstrating that *BIRC6-AS1*/BIRC6 mainly affects the activity of the NHEJ repair pathway induced by abiraterone, thereby promoting the accumulation of DNA damage and apoptosis, and enhancing the resistance of PCa cells to abiraterone (**Figure 10**).

**Figure 10.**
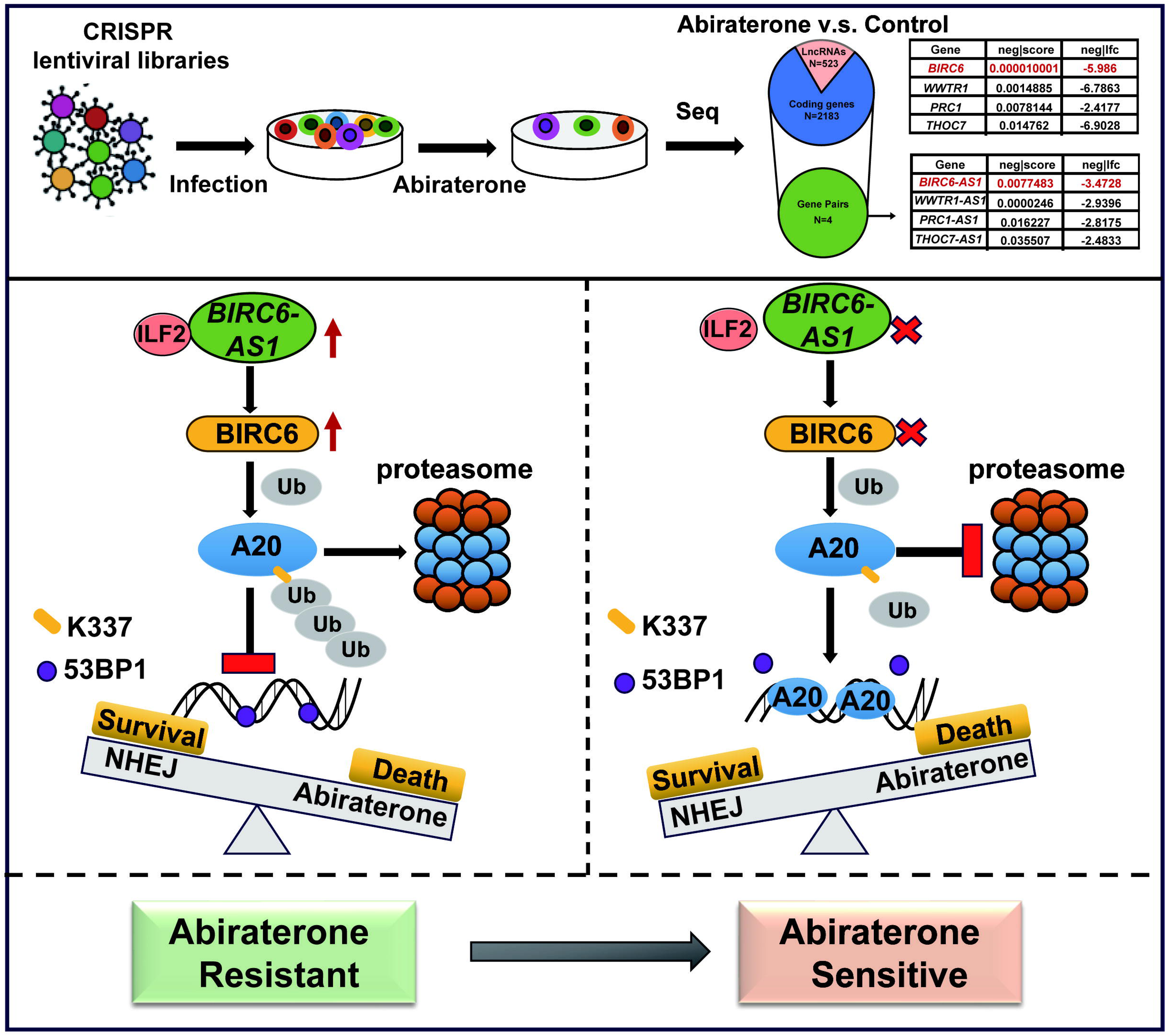
Proposed model for genome-wide CRISPR Cas9 library screening identified *BIRC6-AS1*/BIRC6 as a critical driver for abiraterone resistance in Prostate cancer. *BIRC6-AS1* positively regulates BIRC6 expression by stabilizing its mRNA by interacting with ILF2. Mechanistically, BIRC6 facilitated the ubiquitination and degradation of A20 by specifically targeting lysine 337 (K337), enhanced NHEJ repair activity, leading to abiraterone resistance (abiraterone resistant). *BIRC6-AS1* depletion inhibited BIRC6 expression and prevented A20 degradation, predominantly disrupts NHEJ repair pathway, resulting in the disassembly of 53BP1 foci at DNA damage sites and increased DNA damage accumulation in prostate cancer cells treated with abiraterone (abiraterone sensitive).

Abiraterone, a novel endocrine therapeutic agent, holds a significant place in the treatment paradigm of prostate cancer, particularly in the management of mCRPC. The primary mechanism of action of abiraterone involves the inhibition of the CYP17 enzyme, which disrupts the biosynthesis of androgens, consequently reducing androgen levels and retarding the progression of prostate cancer^20,21^. However, advancing research has unveiled a broader spectrum of abiraterone’s therapeutic effects, transcending mere androgen suppression. Several studies have indicated that specific DDR gene mutations, such as those in BRCA1/2 and ATM, may correlate with sensitivity to abiraterone treatment, with these patients potentially exhibiting superior treatment responses and prognosis ^22,23^. In the context of mCRPC patients who experience therapeutic failure with abiraterone and possess DDR deficiencies, combination therapy employing PARP inhibitors (e.g., olaparib) alongside abiraterone has emerged as a promising strategy ^24,25^. This combinatorial approach aims to further potentiate the cumulative effects on DNA damage, potentially overcoming resistance to abiraterone. Currently, the relationship between abiraterone’s therapeutic mechanism and DNA damage, particularly its interplay with DDR pathways, remains an area of ongoing investigation and is not fully elucidated.

DNA double-strand breaks (DSBs) represent the most severe form of DNA damage, while homologous recombination (HR) repair is one of the crucial DNA repair pathways, with the *BRCA1/2* genes playing a pivotal role. Studies have demonstrated that the activation of the androgen receptor (AR) pathway is associated with the upregulation of DNA damage repair genes, including those involved in HR repair pathway. Abiraterone treatment may impact the AR pathway, thereby modulating the expression of DNA damage repair genes and enhancing the extent of DNA damage ^26,27^. However, there has been limited research on the relationship between abiraterone and NHEJ, another important DNA repair pathway. Our study revealed that abiraterone not only induces HR repair pathway but also triggers NHEJ pathway. In-depth investigation into the correlation between abiraterone and NHEJ can contribute to a better understanding of abiraterone’s therapeutic mechanisms and facilitate the development of more precise and effective treatment regimens for patients with mCRPC.

LncRNAs are involved in regulating multiple biological processes, including proliferation, apoptosis, DNA damage repair and chemotherapeutic resistance^28–34^. For example, lncRNA-p21 is upregulated in neuroendocrine prostate cancer (NEPC) by enzalutamide (Enz) treatment, promoting neuroendocrine differentiation (NED) via altering AR-driven EZH2 function to methylate STAT3^35^. lncRNA *LOC644656* promotes cancer chemoresistance and poor survival by suppressing stemness and modulating DNA damage responses to enhance cell survival under genotoxic stress^36^. Moreover, Damage-induced long non-coding RNAs (dilncRNAs) are transcribed at DSBs, and facilitate DNA damage response (DDR) activation by recruiting repair factors to specific genomic sites^37^. In our study, we found that the expression levels of *BIRC6-AS1* are upregulated in tumor samples compared to paired normal tissues. *BIRC6-AS1* depletion inhibited PCa cells proliferation, and enhanced the sensitivity of PCa cells to abiraterone by inducing cell apoptosis and weakened cell clonal formation ability. The results suggested that *BIRC6-AS1* suppression augments chemosensitivity of PCa cells to abiraterone.

LncRNAs regulate gene expression through multiple pathways, including direct interaction with proteins, interfering with transcription factors binding to target genes, and acting as regulators of transcription factors or chromatin modifiers, among others^11,34,38–40^. Studies have shown that lncRNA *HOTAIR* binds to the AR protein to obstruct its interaction with the E3 ubiquitin ligase MDM2, thereby inhibiting AR ubiquitination and protein degradation^41^. lncRNA *PCBP1-AS1* promoted the deubiquitination of AR/AR-V7 by stabilizing the USP22-AR/AR-V7 complex, thereby inhibiting degradation of AR/AR-V7^42^. LncRNA *NEAT1* drives prostate cancer growth by promoting AGRN expression through its interaction with CDC5L, thereby enhancing proliferation and suppressing DNA damage responses^43^. To determine the underlying mechanism of highly expressed *BIRC6-AS1* enhancing PCa cells resistant to abiraterone, we performed RNA pull-down assay and identified the binding protein ILF2 of *BIRC6-AS1*. The interaction between *BIRC6-AS1* and ILF2 was also verified. We found that inhibition of *BIRC6-AS1* or ILF2 reduced the mRNA and protein levels of BIRC6. Our results also revealed that *BIRC6-AS1* impacts the stability of *BIRC6* mRNA in DU145 cells and PC3 cells by interacting with ILF2 following treatment with actinomycin D. However, how *BIRC6-AS1* modulates the stability of *BIRC6* mRNA needs further investigation.

BIRC6 plays a crucial role in diverse cellular processes, including cell division and the regulation of autophagy ^44^. It is well-recognized for its anti-apoptotic activity. BIRC6 interacts with and inhibits caspases such as caspase-3, caspase-7, and caspase-9, interactions that are themselves inhibited by Smac (second mitochondrial activator of caspases) ^45^. Structurally, BIRC6 is a large multi-domain protein featuring a BIR (Baculoviral IAP Repeat) domain near the N-terminus and an E2-E3 hybrid ubiquitin-conjugating (UBC) domain near the C-terminus ^46,47^. The significance of BIRC6 in cancer biology is highlighted by its overexpression in numerous malignancies, including breast cancer, colorectal cancer and ovarian cancer ^48–50^. Studies have also demonstrated that BIRC6 promotes PCa cell proliferation and impacts the sensitivity of PCa cells to Enzalutamide^16^. To date, the involvement of BIRC6 in abiraterone resistance and the NHEJ process has not been reported. In this study, we found that BIRC6 are highly expressed in tumor samples compared to paired normal tissues. BIRC6 depletion inhibited PCa cells proliferation, and enhanced the sensitivity of PCa cells to abiraterone. We further confirmed that *BIRC6-AS1*/BIRC6 affects the NHEJ pathway when PCa cells are exposed to abiraterone. By modulating the NHEJ pathway, *BIRC6-AS1*/BIRC6 knockout may promote the accumulation of DNA damage and apoptosis, displayed the increased γH2AX levels, thereby enhancing the chemosensitivity of PCa cells to abiraterone, both in vitro and in vivo.

To study the downstream molecular mechanisms by which *BIRC6-AS1*/BIRC6 regulate abiraterone resistance, we predicted potential BIRC6-interacting proteins. Among these three candidates, A20 has been previously reported to suppress NHEJ activity and is essential for the disassembly of 53BP1 at DNA damage foci^19^. Knockout of either *BIRC6-AS1* or BIRC6 enhanced the nuclear localization of A20, and this effect was further potentiated in PCa cells exposed to abiraterone. Given that the BIRC6 protein contains an ubiquitin-conjugating enzyme E2-E3 domain, we hypothesized that BIRC6 may regulate the NHEJ repair pathway through ubiquitination of its substrate. Then, we verified that the UBC domain of BIRC6 was essential for A20 binding, and the 264-548 aa domain of A20 was essential for the binding of BIRC6. Our results also revealed that BIRC6 promoted the protein degradation of A20 depending on the interaction of BIRC6 with A20.

Polyubiquitylation is typically viewed as a posttranslational modification that modulates protein stability or interactions, with linkage types ultimately shaping substrate fate. Studies have shown that The K48-linked polyubiquitin chain serves as the primary degradation signal in eukaryotic cells, targeting modified proteins for recognition and proteolysis by the 26S proteasome ^51,52^. K63-linked polyubiquitin chains bound DNA to recruit repair factors, facilitating DNA damage repair^53^. TRIM21 was reported to mediate K63-linked ubiquitination of RBM38c, thereby suppressing DNA damage-induced autophagy activation^54^. Moreover, FANCG K63-ubiquitination bridges Rap80-BRCA1 for DNA interstrand crosslinks (ICLs)-directed HR repair^55^. Our findings revealed that BIRC6 promotes K48 and K63-linked polyubiquitination of A20. The results indicated that BIRC6 facilitates degradation of A20 via the K48-linked ubiquitin chain, while BIRC6-mediated K63-linked polyubiquitination may correlate with DNA damage repair. However, whether the K63-linked polyubiquitination mediated by BIRC6 is involve in NHEJ repair requires further investigation.

The A20 protein, encoded by the *TNFAIP3* gene, features zinc finger motifs at its carboxyl terminus and an ovarian tumor (OTU) motif at the amino terminus. The A20 protein plays pivotal roles in both biological and clinical contexts, particularly in immune regulation, inflammatory responses, and tumorigenesis and development^56–58^. A20/TNFAIP3 has been identified as a tumor suppressor gene in regulating the progression and metastatic potential of prostate cancer^59^, the findings that align with our research outcomes. Yang et al^19^ identified A20/TNFAIP3 as a novel DNA damage response (DDR) regulator. DNA damage induces NF-κB-mediated A20 transcription, elevating its protein levels at damage sites. A20 inhibits H2A K13/15 ubiquitination by disrupting RNF168-H2A binding, thereby reducing 53BP1 recruitment. This shifts DNA repair balance from NHEJ to HR, optimizing dynamic repair in tumor cells. Conversely, A20 deficiency promotes persistent RNF168/53BP1 accumulation, enhancing NHEJ at the expense of HR efficiency. In our study, the knockout of *BIRC6-AS1* or BIRC6 led to the increased nuclear localization of A20 and facilitated the dissociation of 53BP1 at the site of DNA damage, which was rescued by the inhibition of A20, both in vitro and in vivo. These findings align with the established role of A20 in the NHEJ repair pathway, further reinforcing the notion that BIRC6 regulates the NHEJ pathway via promoting the degradation of A20.

In summary, we conducted a systematic screening for lncRNAs and protein-coding genes associated with abiraterone resistance, confirming positive correlations between abiraterone resistance and DNA repair processes. We unveiled the resistance-mediating function of the paired sense-antisense transcript *BIRC6-AS1*/*BIRC6*, and dissected the regulatory mechanisms involving *BIRC6-AS1*/BIRC6, A20 and the NHEJ repair pathway. Our study provides a comprehensive understanding of the genetic basis of abiraterone resistance and proposes a promising potential target for the treatment of abiraterone-resistant prostate cancer.

## Method

### Plasmids Reagents and antibodies

BIRC6 Antibody (Proteintech Group, Inc, Cat# 29760-1-AP) / (Abcam, Cat# ab19609), A20 Antibody (Santa Cruz Biotechnology, Inc, Cat# sc-166692), ILF2 Antibody (MedChemExpress, Cat# HY-P82199), Flag-tag (Sigma-Aldrich, Cat# F3165), Ub (ABclonal, Cat# A19686), GAPDH (Proteintech Group, Inc, Cat# 60004), Goat anti-Mouse IgG (H+L) HRP Secondary Antibody (Invitrogen, Cat# 31430), Anti-mouse IgG for IP (HRP) (Abcam, Cat# ab131368), Goat anti-Rabbit IgG (H+L) HRP Secondary Antibody (Invitrogen, Cat# 31460), γH2AX Antibody (Servicebio, Cat# GB111841-50), 53BP1 Antibody (HUABIO, Cat# ET1704-05), HA-tag (Beyotime, Cat# AF0039), Abiraterone (Targetmol, Cat# T6216), MG-132 (Targetmol, Cat# T2154), Cycloheximide (Targetmol, T1225).

### Cell culture

HEK-293T and prostate cancer cell lines (DU145 and PC3) were obtained from the American Type Culture Collection (ATCC). All cells were cultured in medium supplemented with 10% fetal bovine serum (ExCell, Cat# FSP500) and 1% penicillin-streptomycin (Beyotime, Cat# C0222), and maintained at 37°C in a 5% CO□ atmosphere. Specifically, HEK-293T and DU145 cells were cultured in Dulbecco’s Modified Eagle Medium (Thermo Fisher Scientific, Cat#12800017), PC3 cells were cultured in RPMI 1640 Medium (Thermo Fisher Scientific, Cat# 31800022). All cell lines were verified to be mycoplasma-free in our study. The prostate cancer cell lines were authenticated by the commercial supplier using microsatellite (short tandem repeat, STR) profiling.

### Preparation of Cas9 expression clones

PC3 cells were transduced with Cas9 lentiviral vector pKLV2-EF1a-Cas9-T2A-Bsd-W (Addgene, Cat# 68343) supplemented with 8 μg/ml polybrene (Solarbio Life Science, Cat# H8761). After 48 h, cells were selected with 15 μg/ml blasticidin (Solarbio Life Science, Cat# B9300) for 4 days to establish PC3 cell lines stably overexpressing Cas9.

50 PC3-Cas9 cells were seeded into 10-cm cell culture dishes and cultured in an incubator. Monoclonal colonies formed after 3-4 weeks, and clones exhibiting robust growth were selected based on morphological observation. The selected clones were trypsinized, transferred to 96-well plates, and expanded to establish cell lines. Cell pellets from monoclonal cultures were collected, and the expression levels of Flag-tagged-Cas9 were detected by Western blotting.

Following successful establishment of clones, Cas9 cleavage efficiency was evaluated using a reporter gene assay in 6-well plates. Cas9 clones were transduced with either pKLV2-U6gRNA5(Empty)-PGK-BFP-2A-GFP-W (Addgene, Cat# 67979) or pKLV2-U6gRNA5(gGFP)-PGK-BFP-2A-GFP-W (Addgene, Cat# 67980). After 3 days, cells expressed both blue fluorescent protein (BFP) and green fluorescent protein (GFP) (control), pKLV2-U6gRNA5(gGFP)-PGK-BFP-2A-GFP-W contained gRNA targeting GFP. Cas9 activity was determined by analyzing the fluorescence expression ratio of GFP to BFP via flow cytometry, and monoclonal cells with the highest activity were selected for subsequent screening.

Cas9 efficiency (%) was calculated as: 100 − [(percentage of GFP□BFP□ cells / (percentage of GFP□BFP□ cells + percentage of GFP□BFP□ cells))×100], representing 100 minus (percentage of uncleaved cells / total percentage of transfected cells×100).

### Genome-wide CRISPR screening

PC3-Cas9 cells (4×10^8^) were transduced with either the Splicing-targeting CRISPR-Cas9 library for human lncRNAs (Addgene, Cat# 119977) or the Human genome-wide lentiviral CRISPR gRNA library version 1 (Addgene, Cat# 67989) at a multiplicity of infection (MOI) of 0.3, ensuring single gRNA integration per cell. After 3 days, cells were selected with 1.5 μg / ml puromycin for 5 days to maximize viral integration and transgene expression. At this stage, 1×10^8^ cells were harvested as baseline controls for next-generation sequencing (NGS). The remaining 1×10^9^ cells were divided into two groups and treated with either DMSO (vehicle control) or abiraterone for 28 days. After treatment, 1×10^8^ cells were harvested per group for analysis. Cell pellets were collected, and genomic DNA was extracted using a GeneJET PCR Purification Kit (Thermo Fisher Scientific, Cat# K0702). sgRNAs were amplified using a two-step PCR strategy, and purified amplicons were analyzed by NGS on the Illumina HiSeq 2500 platform. Primer sequences are provided in Supplementary Table 3.

### Plasmid construction and transfection

The full-length cDNA sequence of *BIRC6-AS1* was synthesized by Beijing Tsingke Biotech Co., Ltd. Subsequently, cDNAs encoding *A20* and *BIRC6-AS1* were subcloned into pcDNA3.1-Myc-His A and pLVX-mCNV-ZsGreen1-Puro vectors, respectively. Using the full-length pLVX-mCNV-ZsGreen1-Puro-BIRC6-AS1 plasmid as template, functional domain deletions of *BIRC6-AS1* were constructed based on predicted secondary structures. Similarly, domain deletions and point mutations of *BIRC6* and *A20* were generated using pDARMO-CMVT-3FLAG-BIRC6 and pcDNA3.1-Myc-His-A20, respectively.

Target gene downregulation was achieved via sgRNAs or shRNAs. sgRNAs targeting *BIRC6*, *BIRC6-AS1*, *RP11 - 1079K10.3*, *WWTR1 - AS1*, *RP11 - 49K24.4*, *WDR18*, *ARFRP1* and *TINF2* were derived from CRISPR sgRNA libraries and cloned into the lentiCRISPRv2 vector. shRNAs against *ILF2* and *A20* were designed and cloned into pLKO.1. For lentivirus production, HEK293T cells were co-transfected with respective shRNA or overexpression constructs (1.5 μg each), envelope plasmid pMD2G (0.375 μg), and packaging plasmid psPAX2 (1.125 μg). All transfections employed Lipofectamine3000 (Invitrogen, Cat# L3000015) or PEI (MedChemExpress, Cat# HY-K2014) according to manufacturer protocols. Primer sequences are provided in Supplementary Table 3.

### Quantitative real-time PCR

Total RNA was extracted from cell lines using TRIzol reagent (Takara, Cat# 9109). RNA was reverse-transcribed into cDNA using the ABScript II cDNA First Strand Synthesis Kit (ABclonal, Cat# RK20400). Quantitative real-time PCR (qRT-PCR) was performed with SYBR Green qPCR Mix (TargetMol, Cat# C0006) according to the manufacturer’s instructions. β*-actin* served as the internal control. Primer sequences are provided in Supplementary Table 3.

### Colony formation assay

For colony formation assays, 1,000 cells were seeded per well in 24-well plates and treated with abiraterone at varying concentrations. After 14 days of incubation, cells were fixed with 4% paraformaldehyde (Sigma-Aldrich, Cat# P6148) and stained with 0.1% crystal violet (Yuanye Bio-Technology Co, Cat# R20756). The number of colonies was determined by counting stained colonies using ImageJ software.

### Cell Proliferation and IC50 Assay

For cell proliferation assay, DU145 or PC3 cells were seeded into 96-well plates at a density of 1×10^3^ cells per well. For IC50 determination, PC3 or DU145 cells were seeded into 96-well plates at a density of 3×10^3^ cells per well and treated with abiraterone at different concentrations.10 μL/well CCK-8 regent (TargetMol, Cat# C0005) was added for 4 h before the indicated time points, and then absorbance was measured at 450 nm using a microplate reader. Values were obtained from quadruplicate wells.

### Apoptosis assay

PC3 and DU145 cell lines were cultured in medium containing abiraterone or vehicle control. After 48 h, cells were harvested by centrifugation and washed twice with ice-cold PBS to remove residual medium and cellular debris. Apoptosis was evaluated using a PE-Annexin V Apoptosis Detection Kit (BIOSCIENCE, Cat# A6030L) following the manufacturer’s protocol. Data were analyzed with FlowJo software (v10.8.1).

### Western blot

Prostate cancer cells were collected and lysed in ice-cold lysis buffer (500 mM NaCl [Yuanye Bio-Technology Co, Cat# R21092]; 20 mM Tris-HCl, pH 8.0 [Yuanye Bio-Technology Co, Cat# R21104]; 1% NP-40 [Solarbio Life Science, Cat# N8030]; 5 mM EDTA [Solarbio Life Science, Cat# E1170]; 1 mM DTT [Solarbio Life Science, Cat# D8220]; 1×Protease Inhibitor [Targetmol, Cat# C0001]) for total protein extraction. Protein concentrations were determined using a BCA Protein Assay Kit (Thermo Fisher Scientific, Cat# 23227). Subsequently, 10-40 μg of protein lysates were separated by 6-12 % SDS-PAGE and transferred to PVDF membranes (Pall Life Sciences, Cat# BSP0161). Membranes were incubated with specific primary antibodies followed by horseradish peroxidase (HRP)-conjugated secondary antibodies. Protein bands were visualized by enhanced chemiluminescence using an ECL kit (BIO-RAD, Cat# 1705061) on X-ray film under darkroom conditions.

### Fluorescence In Situ Hybridization (FISH)

Prostate cancer cells were seeded onto sterile round coverslips in 12-well plates. After fixation with 4% paraformaldehyde, cells were permeabilized at 4 °C for 40 min using 0.2 % Triton X-100 (Sigma-Aldrich, Cat#9036-19-5) in PBS. Subsequently, cells were incubated in 2×SSC solution at 37 °C for 30 min, dehydrated through a graded ethanol series (70%, 85%, and absolute ethanol), and air-dried at room temperature for 15 min. The probe mixture (containing Cy3-labeled probe, pre-incubated hybridization buffer, and DEPC-treated water) was denatured at 73 °C for 5 min, added to each well, and hybridized overnight at 37°C in the dark. The next day, cells were sequentially washed with 0.4×SSC and 2×SSC preheated to 65°C. Subsequently, cells were stained with DAPI at room temperature for 15 min. Finally, coverslips were mounted onto glass slides with 50% glycerol mounting medium and imaged under a fluorescence microscope using a 60×oil-immersion objective. Primer sequences are provided in Supplementary Table 3.

### Nuclear-Cytoplasmic RNA Separation

A total of 5×10^6^ prostate cancer cells were collected. Cytoplasmic and nuclear RNA were isolated using the NE-PER® Nuclear and Cytoplasmic Extraction Reagents (Thermo Fisher Scientific, Cat# 78833) according to the manufacturer’s instructions. Reverse transcription of both RNA fractions was performed using the Reverse Transcription Kit (ABclonal, Cat# RK20400). qPCR was carried out with SYBR Green Master Mix (TargetMol, Cat# C0006), using *U6* and β*-actin* as nuclear and cytoplasmic reference genes, respectively.

### mRNA stability assay

PC3 and DU145 cells (3×10^5^ per well) were seeded in 6-well plates and treated with 10 μg / mL actinomycin D (Titan, Cat# 015005616) for 0, 1, 2, 4, or 8 h. Total RNA was extracted, and *BIRC6* and β*-actin* mRNA levels were quantified by RT-qPCR. mRNA half-lives were calculated by fitting decay curves to mRNA levels over time.

### NHEJ and HR reporter assays

NHEJ and HR reporter assays were performed according to previously published methods^60,61^. Briefly, The NHEJ reporter plasmid pEJ5-GFP (Addgene, Cat# 44026) or HR reporter plasmid DR-GFP (Addgene, Cat# 26475) was transfected into DU145 and PC3 cells, followed by selection of stable clones with 1 μg /ml puromycin. To induce DNA double-strand breaks (DSBs), lentiviruses encoding sgBIRC6, sg*BIRC6-AS1*, or control sgRNA were transduced into DU145 and PC3 cells stably expressing pEJ5-GFP or DR-GFP, respectively. Concurrently, the pCBASce plasmid (Addgene, Cat# #26477) was transfected into these cells under conditions with or without abiraterone treatment. Finally, GFP-positive cells were quantified by flow cytometry.

### Immunofluorescence

DU145 or PC3 cells (1.5×10^4^) were seeded on coverslips in 24-well plates. After 24 h, cells were treated with 30 μM abiraterone or vehicle for 48 h. Cells were washed with PBS, fixed in 4% PFA (15 min, RT), permeabilized with 0.2% Triton X-100 (15 min), and blocked in 5% BSA (1 h, 37°C) (Sigma-Aldrich, Cat# B2064). Primary antibodies (anti-γH2AX, anti-53BP1, anti-A20) diluted in blocking buffer were applied overnight at 4°C. After PBS washes, Alexa Fluor 488/594-secondary antibodies (1:500) were incubated for 1 h (RT, dark). Nuclei were stained with DAPI (0.5 μg/ml, 15 min). Stained cells were imaged using a high-resolution laser confocal microscope or upright fluorescence microscope.

### Xenograft models

All animal procedures were approved by the Animal Ethics and Welfare Committee of Shaanxi Normal University (Approval No. 2024-076). Tumor volumes did not exceed the committee’s maximum permitted size (2000 mm^3^). Four-week-old male BALB/c nude mice (BALB/c-nu/nu; Beijing HFK Bioscience Co, Ltd.) were housed under specific pathogen-free (SPF) conditions with a 12-h light / dark cycle, ad libitum access to water and standard chow. After one-week acclimatization, mice were randomized into groups (n = 5).

Prostate cancer cells were resuspended in ice-cold PBS and counted. Single-cell suspensions (150 μL) containing 1×10^6^ PC3 or 3.5×10^6^ DU145 cells were injected subcutaneously into the right flank. When tumors reached ∼50 mm^3^, mice were re-randomized into treatment groups. Abiraterone was dissolved in DMSO and diluted in corn oil. Mice received abiraterone (150 mg/kg) or vehicle control via oral gavage every other day. Tumor dimensions were measured every 3 days with digital calipers, and volumes calculated as (Length×Width^2^) ×0.5. At day 20 post-treatment, mice were euthanized, and tumors were excised and fixed in 4% paraformaldehyde for histological analysis.

### RNA Pull-down

PCR products amplified from the pLVX-mCNV-ZsGreen1-Puro-BIRC6-AS1 template, including full-length *BIRC6-AS1* and its truncation variants, were separated on 1.5% agarose gels and validated by electrophoresis. Target bands were excised and purified. Biotinylated RNA was synthesized via in vitro transcription of these templates using the High Yield T7 Biotin16 RNA Labeling Kit (APExBIO, Cat# K1082) with Biotin-16-UTP, followed by DNase I treatment (1 U/μL, 37°C, 15 min) to digest residual DNA. Purified RNA was heat-denatured (90 °C, 2 min), snap-cooled on ice (5 min), and refolded at 25 °C (30 min) to establish secondary structures. Primer sequences are provided in Supplementary Table 3.

DU145 cells were lysed in ice-cold RIP buffer (50 mM Tris-HCl [pH 7.4], 150 mM NaCl, 1% NP-40, 0.5% Sodium deoxycholate [Sigma-Aldrich, Cat# D6750], 0.1% SDS [Invitrogen, Cat# 15553], protease inhibitor and RNase inhibitor [Applied Biosystems, Cat# N8080119]) for 30 min on ice. Lysates were centrifuged (14,000×g, 15min, 4 °C), and supernatants were collected with 10% reserved as input. Streptavidin Agarose (Beyotime, Cat# P2159) were washed thrice with RIP buffer and incubated with denatured biotinylated RNA (3 μg) at 25 °C for 1 h with rotation. After three RIP buffer washes, beads were incubated with cell lysate at 4 °C for 2 h to capture protein-RNA complexes. Complexes were eluted in 2×SDS loading buffer supplemented with 100 mM DTT and 10 mM biotin, then resolved by SDS-PAGE for Western blotting or prepared for LC-MS/MS analysis.

### Mass Spectrometry Analysis of RNA Pull-Down Products

Following RNA pull-down, equal amounts of samples pulled down by sense and antisense *BIRC6-AS1* were resolved via 12% SDS-PAGE and visualized using a Fast Silver Stain Kit (Beyotime, Cat# P0017S). Specific protein bands were excised and subjected to LC-MS/MS analysis by a commercial biotech company. Protein identification was performed against the human RefSeq protein database (National Center for Biotechnology Information) using Mascot v2.4.01.

### Comet assay

Prostate cancer cells were treated with or without abiraterone for 48 h, then resuspended in pre-cooled PBS at a concentration of 1×10^6^ cells/mL. DNA double-strand breaks (DSBs) were evaluated using a Comet Assay Kit (Beyotime, Cat# C2041M) according to the manufacturer’s instructions. Briefly, fresh cell suspensions were mixed with 0.7% low-melting-point agarose, spread onto comet slides pre-coated with 1% normal-melting-point agarose, and solidified at 4 °C for 10 min. The slides were then lysed in lysis buffer for 1-2 h, followed by DNA unwinding in alkaline buffer (pH >13) for 60 min. Electrophoresis was conducted at 25 V for 20-30 min in the dark using pre-cooled electrophoresis buffer. After neutralization (performed 1-3 times for 5-10 min each at 4 °C), the slides were stained with 20 μL Propidium Iodide (PI) in the dark for 10–20 min, washed with ultrapure water, covered with coverslips, and observed under an upright fluorescence microscope.

### Co-immunoprecipitation assay

Cells were lysed on ice in lysis buffer (500 mM NaCl; 20 mM Tris-HCl, pH 8.0; 1% NP-40; 5 mM EDTA; 1 mM DTT; 1× Protease Inhibitor) for 30 min. Lysates were centrifuged at 12,000 g for 15 min at 4 °C, and 10% of the supernatant was reserved as input. The remaining lysate was incubated with primary antibody (3 μg) overnight at 4 °C with rotation. Pre-washed Protein A/G agarose beads (Yeasen Biotech, Cat# 36403ES) were added the next day, followed by incubation for 6–8 h at 4°C. Co-immunoprecipitation complex was washed three times with lysis buffer and pelleted by centrifugation at 12,000 g for 5 min at 4 °C. Bound proteins were eluted by boiling in 2×SDS sample buffer and analyzed by SDS-PAGE/immunoblotting.

### Protein half-life assay

HEK-293T cells were seeded in 6-well plates for transient transfection. For proteasome inhibition studies, cells were pretreated with 10 μM MG-132 or DMSO vehicle control for 8 h. For protein stability assays, cells were incubated with 50 μg/mL cycloheximide (CHX) for 0, 12, 18, or 24 h, respectively. Cells were harvested and lysed to extract total protein, followed by Western blot analysis.

### Ubiquitination Assay

Cell pellets were collected, lysed with lysis buffer, and subjected to immunoprecipitation. Ubiquitinated target proteins in the immunoprecipitates were then detected by immunoblotting using an anti-ubiquitin antibody or anti-HA.

### Immunohistochemistry (IHC)

IHC staining of xenografts was performed according to previously published methods^62^. Briefly, 4-μm sections were cut from paraffin-embedded samples, deparaffinized in xylene for 10 min, rehydrated through a graded ethanol series (100–70%), and incubated with antigen retrieval solution followed by 3% H□O□ for 10 min each to block endogenous peroxidase activity. Sections were rinsed with ddH_2_O and incubated overnight at 4 °C with primary antibodies (Anti-BIRC6, Anti-γH2AX, Anti-A20, and Anti-53BP1). Following PBS washes, sections were incubated with horseradish peroxidase (HRP)-conjugated secondary antibodies for 60 min at room temperature, developed with DAB substrate for 5-10 min, and counterstained with hematoxylin. Sections were dehydrated through a graded ethanol series, cleared in xylene, and mounted with neutral balsam. Staining was visualized using a brightfield microscope. IHC scoring was calculated as: (Positive cell percentage score)×(Staining intensity score). Positive cell percentage:0 (no staining), 1 (1-14 %), 2 (15-49 %), 3 (50-74 %), 4 (≥75 %). Staining intensity: 0 (negative), 1 (faint yellow), 2 (light brown), 3 (moderate brown), 4 (dark brown). Tissues were classified as high-expression or low-expression using the cohort median score as threshold.

### Molecular docking study

To evaluate the binding affinity between BIRC6-UBC and A20 proteins, molecular docking was performed using Zdock (https://zdock.wenglab.org/) following the standard protocol. The procedure consisted of the following sequential steps: (1) Retrieval of target protein structures from the Protein Data Bank (PDB.org); (2) Removal of non-essential substructures and water molecules; (3) Addition of hydrogen atoms; (4) Calculation of atomic charges using the Gasteiger method; (5) Execution of 100 independent docking simulations to generate conformational ensembles; (6) Optimization of binding partners’ positional, conformational, and orientational space. The Lamarckian Genetic Algorithm (LGA) was employed for all simulations. Protein-protein interaction maps were generated, and docking outcomes—including binding energy (kcal / mol) and key interacting residues—were systematically recorded.

### Statistics and reproducibility

All statistical analyses were performed using GraphPad Prism v10.1.1 (GraphPad Software, San Diego, CA). Data are presented as mean ± standard deviation (SD). All comparisons were performed using two-way ANOVA or one-way ANOVA, as specified in the figure legends. Statistical significance was defined as *P* < 0.05. For in vitro experiments, at least three biologically independent experiments were performed unless stated specified.

## Data availability

Raw sequencing data of the CRISPR library supporting the findings of this study have been deposited in the National Center for Biotechnology Information (BioProject ID: PRJNA1304892 and PRJNA1306795).Gene expression and clinical information were obtained fromTCGA,ICGC and GSE databases (GSE59745, GSE94767, GSE21034, GSE30521, GSE49244 and GSE85672). The mass spectrometry proteomics data have been deposited to the ProteomeXchange Consortium via the PRIDE^63^ partner repository with the dataset identifier PXD066934.

## Acknowledgements

P.G. was supported by National Natural Science Foundation of China (82472665) and Fundamental Research Funds for the Central Universities (GK202505003), X.M.D. was supported by Natural Science Foundation of Shaanxi Province (2023-JC-YB-716).

## Author Contributions

X.M.D., and P.G. conceived and designed the project. L.L. assistant by X.N.A, Y.T.R., R.Y., P.L., X.Y.W. and X.C.H. performed the experiments. L.L. performed data analysis. X.M.D., P.G. and L.L. wrote the manuscript.

