## Supplementary Figures for "CRlSPR/Cas9 screening revealed *BlRC6-AS1*/BlRC6 mediates abiraterone resistance via NHEJ pathway-dependent A20 degradation in prostate cancer"

**Supplementary Figure 1**

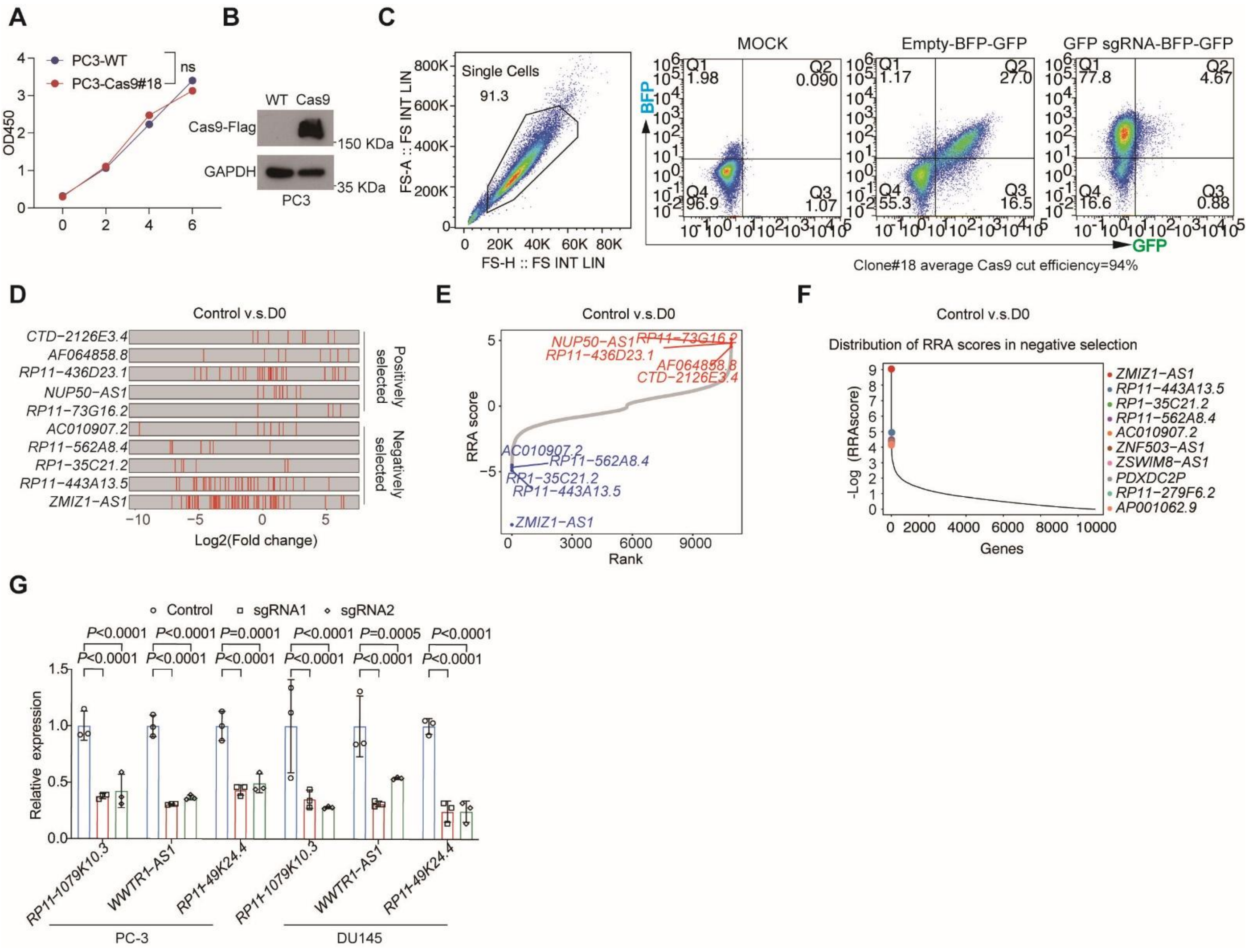


### Figure S1 CRISPR library screening identified lncRNAs association with abiraterone resistance

(**A**) The cell proliferation was measured in Cas9-overexperssion PC3 cells by CCK-8 assay. Data are shown as the mean ± SD (n = 4 biological replicates). (**B**) Western blotting analysis of Cas9 protein level in control and Cas9 overexpressed PC3 cells. GAPDH was used as a loading control. (**C**) Flow cytometer analysis of cleavage efficiency in Cas9-expressing PC3 cells. (**D**) Frequency distribution of $log^2$ fold change for all sgRNAs (top) and $log^2$ fold changes of individual sgRNAs targeting representative candidate genes (bottom), derived from lncRNA CRISPR screens (Control vs. D0). (**E**) The RRA score distribution plot reveals the 10 top-tier candidate essential genes. (**F**) The RRA score distribution plot reveals the top 10 candidate LncRNAs associated with abiraterone resistance. (**G**) RT-qPCR analysis of knockdown efficiency of *RP11-1079K10.3*, *WWTR1-AS1* and *RP11-49K24.4* in PC3 and DU145 cells. Data are mean ± SD (n = 3 biological replicates). Data were analyzed by Ordinary two-way ANOVA（**A**） or two-way ANOVA with Dunnett’s（**G**） multiple comparisons test.

#### Supplementary Figure 2

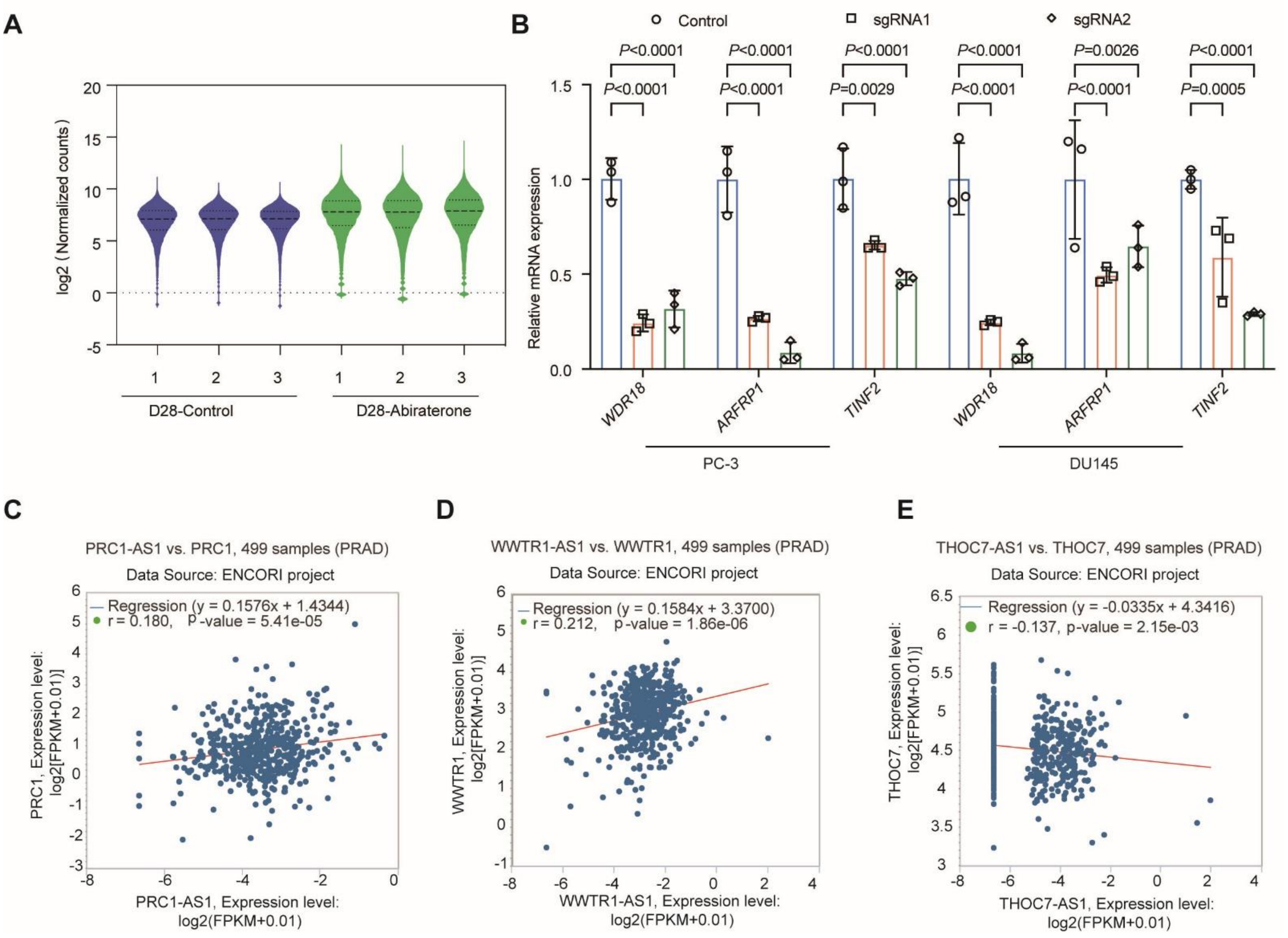


**Figure S2 Co-expression analysis of the three sense-antisense gene pairs that were screened from CRISPR library**

(**A**) Box plots displaying sgRNA distribution in the experimental groups from lncRNA CRISPR-Cas9 library: D28-DMSO (vehicle control) and D28-Abiraterone (treatment). (**B**) RT-qPCR analysis of knockdown efficiency of *WDR18*, *ARFRP1 and TINF2* in PC3 and DU145 cells. Data are shown as the mean ± SD (n = 4 biological replicates). (**C-E**) Co-expression analysis of three sense-antisense gene pairs (*PRC1-AS1* vs *BIRC6*, *WWTR1-AS1* vs *WWTR1* and *THOC7-AS1* vs *THOC7*) in prostate samples was performed based on the ENCORI database. Data were analyzed by two-way ANOVA with Dunnett'smultiple comparisons test（**B**）.

**Supplementary Figure 3**

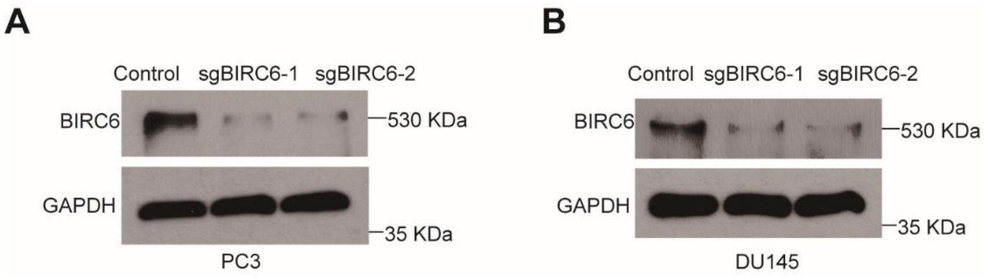


**Figure S3 Western blot analysis of knockdown efficiency of BIRC6 in PCa cells**

(**A** and **B**) Western blot analysis of BIRC6 protein levels in BIRC6 knockdown PC-3 cells (**A**) and DU145 (**B**) cells. GAPDH was used as a loading control. The samples derived from the same experiment and the gels/blots were processed in parallel.

#### Supplementary Figure 4

A

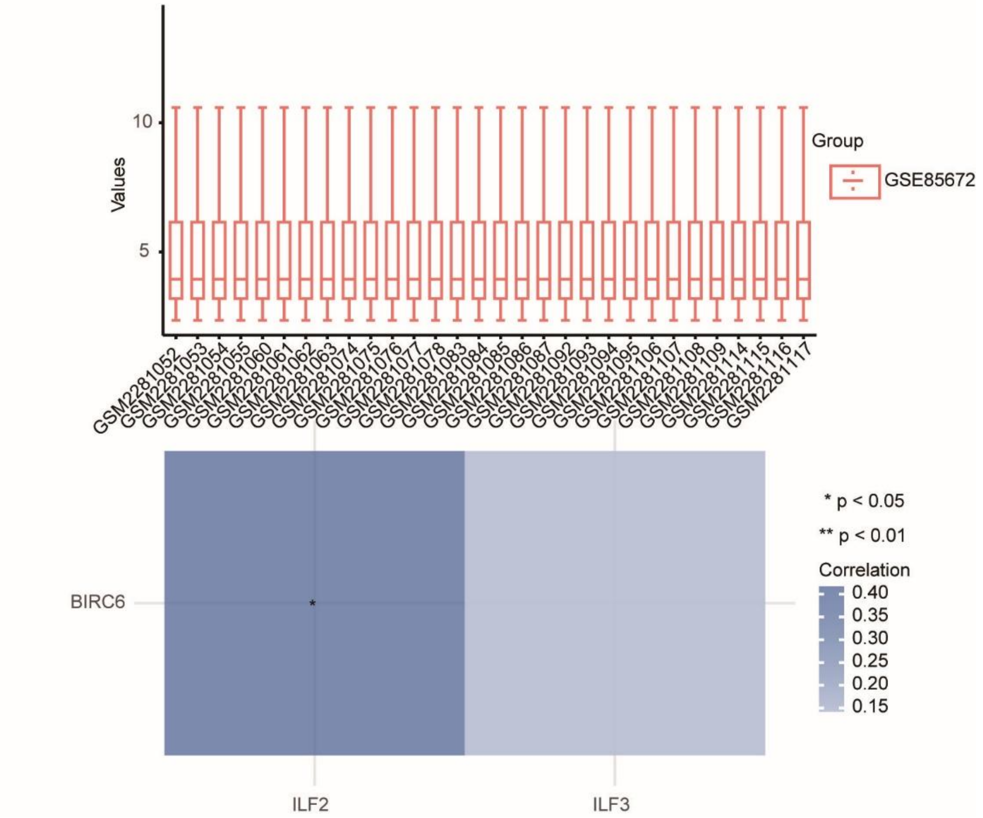


#### Figure S4 The correlation between BIRC6 and ILF2.

(**A**) Co-expression analysis of BIRC6 and ILF2 was performed based on the GSE85672 cohort comprising 30 Abiraterone-treated CRPC prostate tumors.

**Supplementary Figure 5**

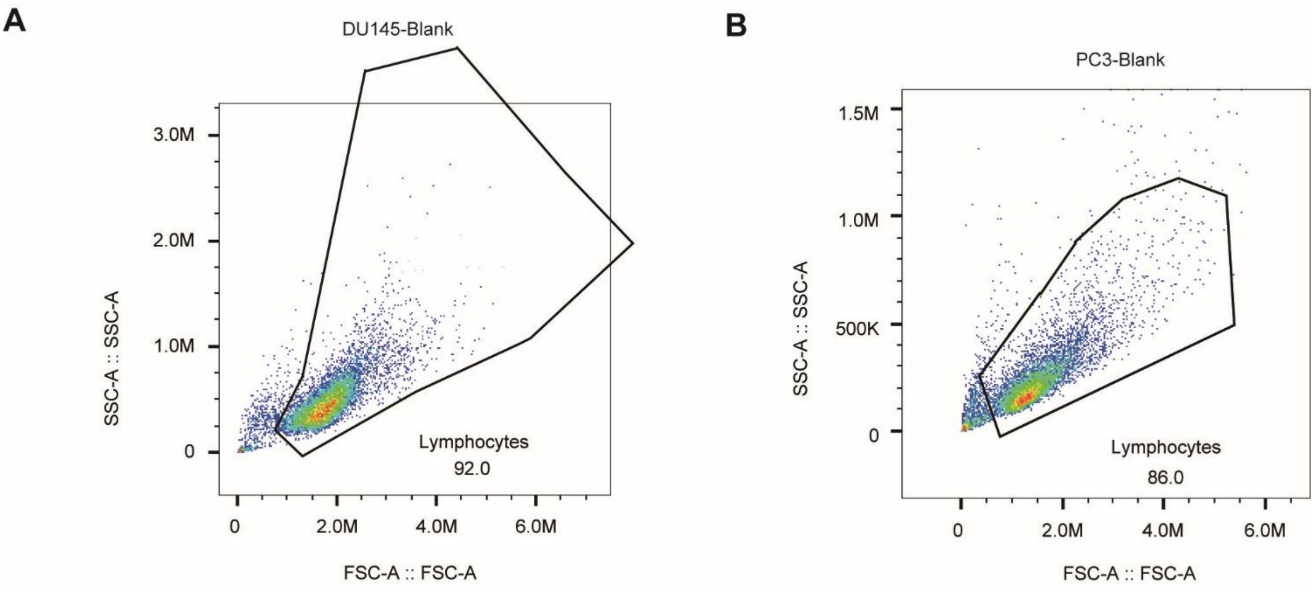


**Figure S5 Flow cytometry scatter plots showing cell population distribution and gating strategy for control samples**

(**A** and **B**) Flow cytometry scatter plots of DU145 untreated control cells (**A**) and PC3 untreated control cells (**B**), showing cell population distribution and gating strategy for control samples.

#### Supplementary Figure 6

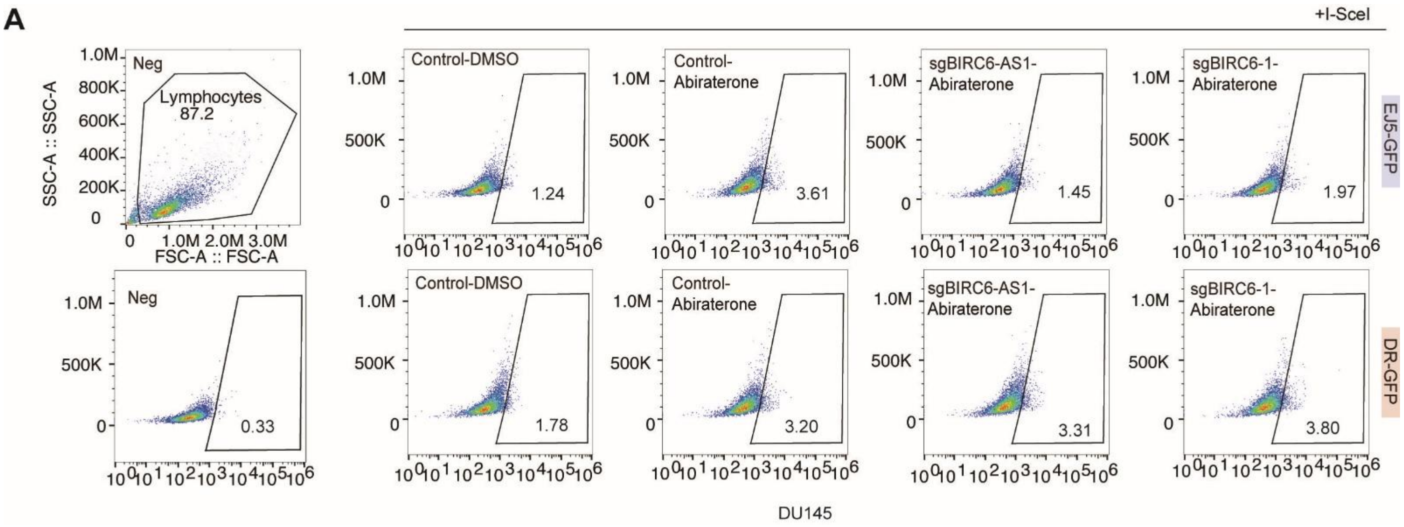


**Figure S6 Flow cytometer analysis of NHEJ and HR repair efficiency in *BIRC6-AS1* or BIRC6 depletion DU145 cells in the presence of abiraterone**

(**A**) Flow cytometer analysis of non-homologous end joining (NHEJ) and homologous recombination (HR) repair efficiency in *BIRC6-AS1* or BIRC6 depletion DU145 cells cultured with or without abiraterone (30 µM).

**Supplementary Figure 7**

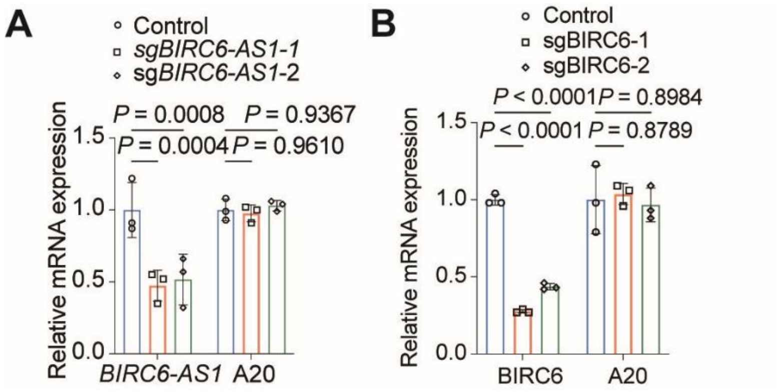


#### Figure S7 qRT-PCR analysis of A20 mRNA levels in *BIRC6-AS1* or BIRC6 depletion DU145 cells

(**A** and **B**) qRT-PCR was used to measure *A20* mRNA levels in DU145 cells transduced with control sgRNAs or sgRNAs targeting *BIRC6-AS1* (**A**) and *BIRC6* (**B**). Data are shown as the mean ± SD (n = 3 biological replicates). Data were analyzed by two-way ANOVA with Dunnett's multiple comparisons test.

**Supplementary Figure 8**

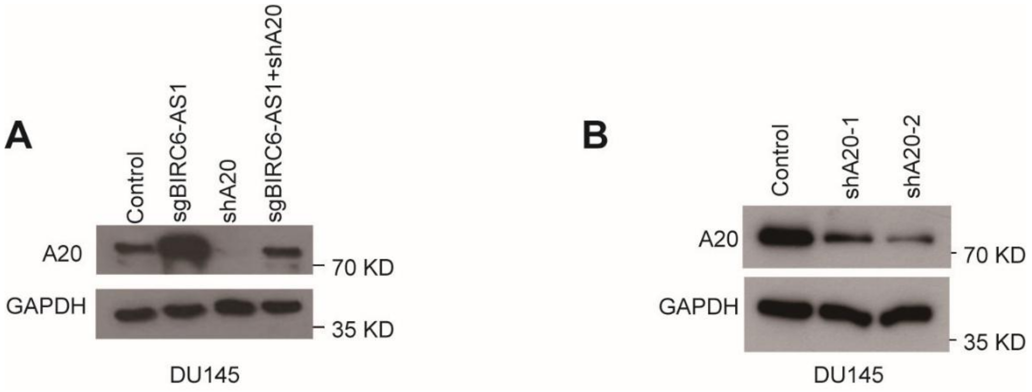


**Figure S8 Western blot analysis of A20 protein levels in various DU145 cell strains**

(**A**) Western blot analysis of A20 protein levels in the indicated DU145 cell strains (Control, sgBIRC6, shA20, and sgBIRC6+shA20 groups). GAPDH was used as a loading control. (**B**) Western blot analysis of A20 protein levels in A20 knockdown DU145 cells. GAPDH was used as a loading control.
